# Protein degradation shapes developmental tempo in mouse and human neural progenitors

**DOI:** 10.1101/2024.08.01.604391

**Authors:** Shota Nakanoh, Despina Stamataki, Lorena Garcia-Perez, Chiara Azzi, Hayley L Carr, Alexandra Pokhilko, Loukik Doshi, Giulia L M Boezio, Manuela Melchionda, Lu Yu, Steven Howell, Mark Skehel, David Oxley, Simon Andrews, James Briscoe, Teresa Rayon

## Abstract

The pace of embryonic development differs markedly across mammalian species, yet the molecular mechanisms underlying these tempo differences remain largely unknown. Here, we systematically compared protein dynamics in mouse and human neural progenitors (NPs) and examined how protein stability influences developmental timing. We find that mouse NPs exhibit faster protein production and degradation than human NPs. Human NPs display broadly increased protein half-lives, independent of cellular compartment or protein function, and this difference persists in post-mitotic neurons. Consistent with this, proteasomal activity is lower in human embryonic spinal cord and stem cell–derived neural progenitors than mouse, correlating with reduced expression of proteasome-associated proteins. Functionally, accelerating the degradation of the key transcriptional repressor IRX3 in mouse NPs speeds the activation of its target gene, providing causal evidence that protein turnover modulates developmental tempo. These results reveal that species-specific regulation of protein degradation shapes the timing of neural development and suggest that evolutionary tuning of proteasomal activity contributes to differences in embryonic developmental pace.

## INTRODUCTION

The developmental mechanisms governing embryogenesis are remarkably conserved across mammalian species, yet the pace at which these programmes unfold is species-specific^1–3^. These species differences in developmental timing are recapitulated *in vitro* using stem cell models, which has allowed investigations into the mechanisms controlling tempo^2–7^. Previous work implicated protein metabolism, the balance of protein synthesis and degradation, as an important contributor to developmental tempo^4,5^. Specifically, reduced rates of protein degradation corresponded to slower tempos in human neural progenitors (NPs) compared to mouse. Similarly, the pace of the segmentation clock during the elongation of the vertebrate axis correlates with the rate of protein turnover in a variety of mammalian species^8^.

While these findings suggest protein turnover is crucial for tempo control, several questions remain unresolved. The regulation of the production and degradation machineries and their dynamics during development are poorly understood. More precisely, a systematic characterization of the similarities and differences in protein production and half-lives between species is lacking, and it remains to be determined whether overall abundance of proteins in embryos differs between species due to species-specific protein turnover dynamics, or compensatory mechanisms adjust protein synthesis rates to achieve equivalent steady-state protein levels. Finally, whether manipulating the stability of individual proteins affects the pace of development is unclear.

Here, we address these questions by quantifying protein production and degradation in equivalent mouse and human NPs. Combining targeted protein labelling, high-resolution quantitative mass spectrometry, proteasomal activity quantifications, and protein depletion with self-labeling tags, we systematically identify species differences in protein production and degradation rates and uncover the underlying mechanisms. Specifically, we elucidate the relative contributions of active protein degradation and dilution via cell division to tempo divergence. Importantly, we demonstrate for the first time that proteasomal activity is lower in human embryonic spinal cords than in stage-matched mouse embryos. Furthermore, we provide evidence to support a role for proteolytic degradation in dictating developmental tempo by decreasing the stability of a key regulatory protein within the gene regulatory network governing motor neuron formation in the neural tube.

Collectively, our work offers a comprehensive comparative analysis of protein dynamics between two mammalian species. We report species-specific differences in protein production and degradation rates and provide *in vivo* evidence that degradative machineries are less active in human embryos than in mice. Our findings highlight the centrality of protein degradation in controlling developmental tempo, and have broad implications for understanding phenotypic diversity across evolution.

## RESULTS

### SNAP Tagged OLIG2 demonstrates differences in protein turnover between mouse and human neural progenitors

The balance between protein synthesis and degradation (protein metabolism) determines protein levels in cells. Having observed global differences in protein decay between mouse and human neural progenitors (NPs)^4^, we set out to test if similar differences were observed for specific proteins. To this end we focused on OLIG2, an essential transcription factor for motor neuron differentiation^9,10^. To obtain data on the turnover for OLIG2 in mouse and human NPs, we tagged the C-terminus of endogenous OLIG2 in mouse and human embryonic stem cells with a HA tag and a SNAP tag, a self-labelling protein tag that enables the covalent labelling by substrates^11^.

We confirmed that the differentiation dynamics of OLIG2::HA::SNAP (OLIG2-SNAP) targeted cells followed the expected pattern for OLIG2 in mouse and human NPs, with OLIG2 increasing upon Retinoic Acid (RA) and Smoothened Agonist addition (SAG)^4^. We detected OLIG2 positive cells at the expected days of differentiation, and there was a good correlation between OLIG2 antibody and HA tag (Fig. S1a). This indicated that tagging does not affect OLIG2 dynamics or function in mouse or human NPs. Next, we tested the specificity of the SNAP fluorescent substrate SNAP 647-SiR. We incubated human NPs from day 4 or day 6 of differentiation with SNAP or HALO ligands conjugated to a 647 fluorophore. We detected specific labelling with the SNAP 647-SiR ligand; the HALO-JF647 substrate did not cross-react with the hOLIG2-SNAP line (Fig. S1b).

Having established ligand specificity, we then determined the half-life of OLIG2 in mouse and human NPs. First, we cultured NPs for 1 hour in media containing SNAP 647-SiR ligand. This was then replaced with media containing SNAP-ligands without 647 fluorophore or SNAP-cell block, a cell permeable and non-fluorescent compound that blocks the reactivity of the SNAP-tag. We observed similar trends in the decay of OLIG2 intensity in human NPs at day 4 and day 6 of differentiation (Fig. S1c), suggesting similar protein decay rates for OLIG2 during differentiation. Comparative SNAP pulse-chase analysis indicated the OLIG2 half-life was ∼1.4-fold longer in human compared to mouse (mOLIG2 t1/2 = 2.31h ± 0.43; hOLIG2 t1/2 = 3.21h ± 0.30) (Fig. 1a). These data corroborate an increased OLIG2 stability in human compared to mouse^4^.

**Figure 1.**
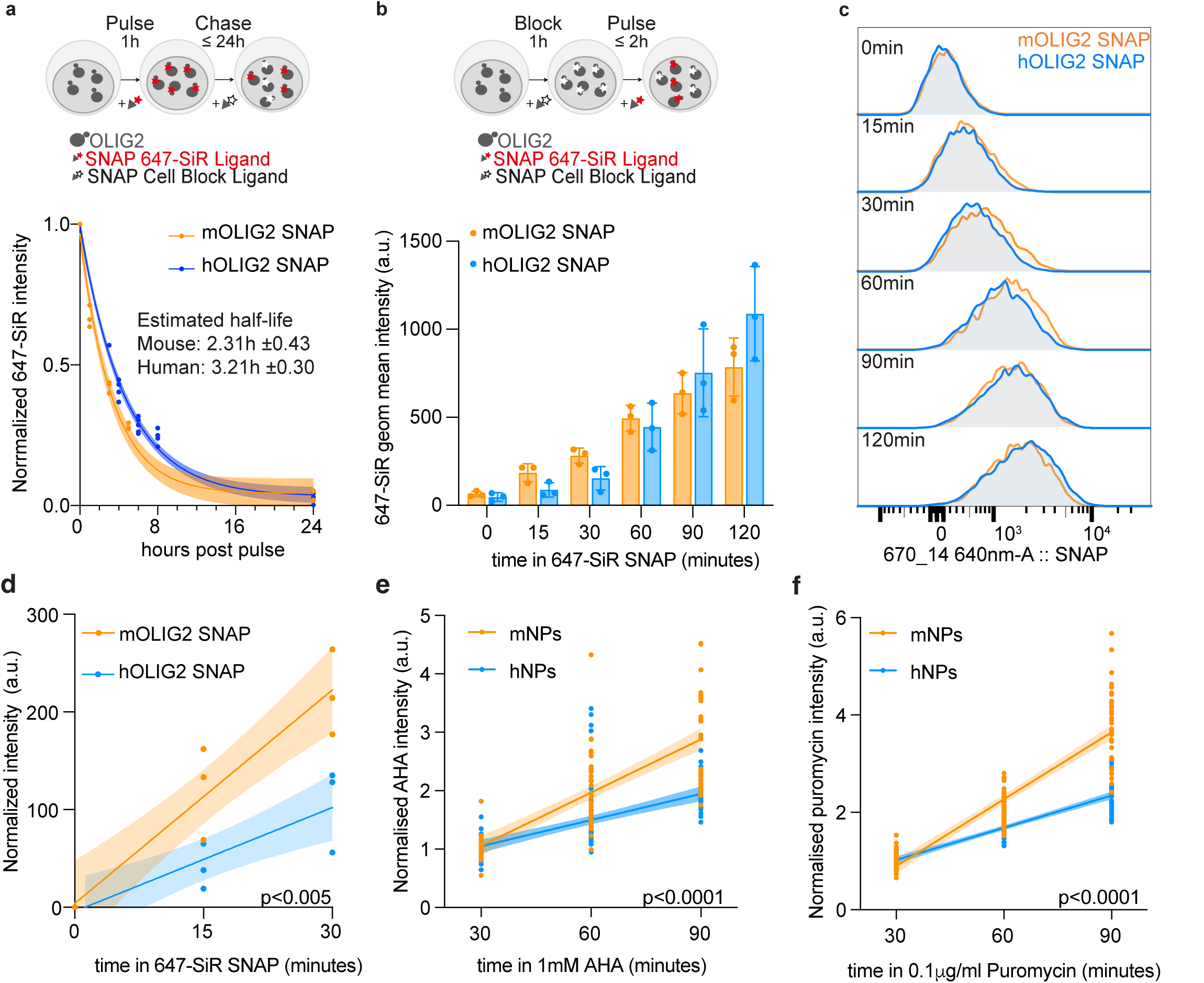
Dynamics of protein production and degradation in mouse and human neural progenitors. (a) OLIG2-SNAP pulse and chase experiments in mouse (orange) and human (blue) NPs. Curves and shaded areas show exponential fit and 95% confidence intervals, respectively (bottom); (mouse n=3, human t_0h_ n=5; t_3h_ n=1; t_4h_=4; t_6h_ n=5; t_8h_ n=4; t_24h_ n=5). (b) OLIG2-SNAP production measurements in mouse and human NPs. Geometric mean ± standard deviation of SNAP 647-SiR ligand incorporation at the indicated time points in mouse and human NPs (n=3). (c) Representative flow cytometry analysis of 647-SiR ligand incorporation in mouse and human NPs for OLIG2-SNAP production measurements. (d) Quantification of OLIG2-SNAP production at the indicated time points in mouse and human NPs. Values correspond to the normalized intensity corrected to 0 minutes, and lines represent a simple linear regression interpolation with 95% confidence intervals. Estimated slopes: mOLIG2 = 7.281 ± 2.26 a.u.·min^-1^; hOLIG2 = 3.544 ± 1.75 a.u.·min^-1^. P-value corresponds to an Analysis of Covariance (ANCOVA) test (n=3). (e) Quantification of AHA incorporation at the indicated time points in mouse and human NPs, normalized to the initial value at 30 minutes. Dots correspond to the geometric mean intensity, and lines represent a simple linear regression interpolation with 95% confidence intervals. Estimated slopes: mNP 0.030 ± 0.0023 a.u.·min^-1^; hNPs 0.015 ± 0.0016 a.u.·min^-1^. Data are shown as n = 4, with at least two technical replicates per experiment. P-value corresponds to ANCOVA test. (f) Quantification of puromycin (0.1 µg/ml) incorporation at the indicated time points in mouse and human NPs normalized to the initial value at 30 minutes. Dots correspond to the geometric mean intensity, and lines represent a simple linear regression interpolation with 95% confidence intervals. Estimated slopes: mNP 0.046 ± 0.0019 a.u.·min^-1^; hNPs 0.022 ± 0.001 a.u.·min^-1^. Data are shown as n = 4, with at least two technical replicates per experiment. P-value corresponds to ANCOVA test.

We next determined OLIG2 production rates for mouse and human. SNAP ligands bind to mature proteins (Fig. 1b). To distinguish between old OLIG2 and newly synthesized molecules, we cultured mouse and human NPs with SNAP-cell block to obstruct all pre-existing OLIG2 molecules present in cells. NPs were then switched into media containing SNAP 647-ligand and samples collected at intervals over the following two hours (Fig. 1b, 1c). Quantifications of the mean intensities for the ligand at various time points indicated that OLIG2 production rates in human NPs were slightly lower in comparison to mouse at 15 min and 30 min (Fig. 1b). Given that intensity measurements will also be affected by protein degradation, and OLIG2 degradation rates are different between mouse and human, we calculated the slope of mouse and human incorporation as a proxy for OLIG2 production rates up to 30 minutes. The accumulation of 647-SIR was ∼2.05-fold steeper in mNPs than in hNPs, raising the possibility of slower protein production rate in human cells (mOLIG2 = 7.281 a.u.·min^-1^ ± 2.26; hOLIG2 = 3.544 a.u.·min^-1^± 1.75) (Fig. 1d).

### Higher protein production rate in mouse versus human neural progenitors

Given the trend for higher OLIG2 production in mouse, we next set out to compare global protein production rates in mouse and human NPs. We performed metabolic labelling of nascent proteins by adding the methionine analog L-azidohomoalanine (AHA)^12^ to media and determined the rate of incorporation over one hour. We treated cells with AHA and collected mouse and human NPs after 30, 60 or 90 minutes of treatment. The production rate was ∼2.3-fold higher in mouse compared to human NPs (mNP = 0.03 a.u.·min^-1^ ±0.005; hNP = 0.014 a.u.·min^-1^ ± 0.003) (Fig. 1e). To confirm higher production rates between mouse and human NPs, we measured puromycin incorporation rates^13^. While both AHA and puromycin incorporate in nascent polypeptides, puromycin blocks protein synthesis and causes premature termination, with the resulting truncated polypeptides often failing to fold properly^14,15^. We treated mouse and human NPs with various concentrations of puromycin to identify the lowest concentration that allowed sufficient labelling for quantification. Puromycin concentrations of 0.1 μg/ml were sufficient to detect nascent protein in mouse and human NPs without triggering a stress response, as measured by levels of EIF2A phosphorylated at Ser51^16^ (Fig. S1d-f). Side-by-side puromycin incorporation assays in mouse and human NPs confirmed a significantly higher ∼2.07-fold rate of protein production in mouse (mNP= 0.045 a.u.·min^-1^ ±0.004; hNP = 0.022 a.u.·min^-1^ ± 0.002) (Fig. 1f). The puromycin results were consistent with the measurements of AHA and confirmed that the rates of protein production are higher in mouse NPs.

### Global proteomic differences associated to protein degradation in mouse and human NPs

Next, we asked if specific sets of proteins showed a greater species difference than others. We performed data-independent acquisition mass spectrometry (DIA-MS) of mouse and human NPs, and calculated protein copy number per cell using the ‘proteomic ruler’ approach^17^ (see Methods) (Fig. S2a-e). We identified 5718 orthologous proteins between mouse and human (Table S1), which we used for differential expression analysis. We found 1791 differentially expressed proteins with ≥ 2-fold change and FDR ≤ 0.05 (Table S2): 997 proteins upregulated in human and 794 in mouse (Fig. 2a). The differentially expressed proteins included lowly and highly abundant proteins (Fig. S2f).

**Figure 2.**
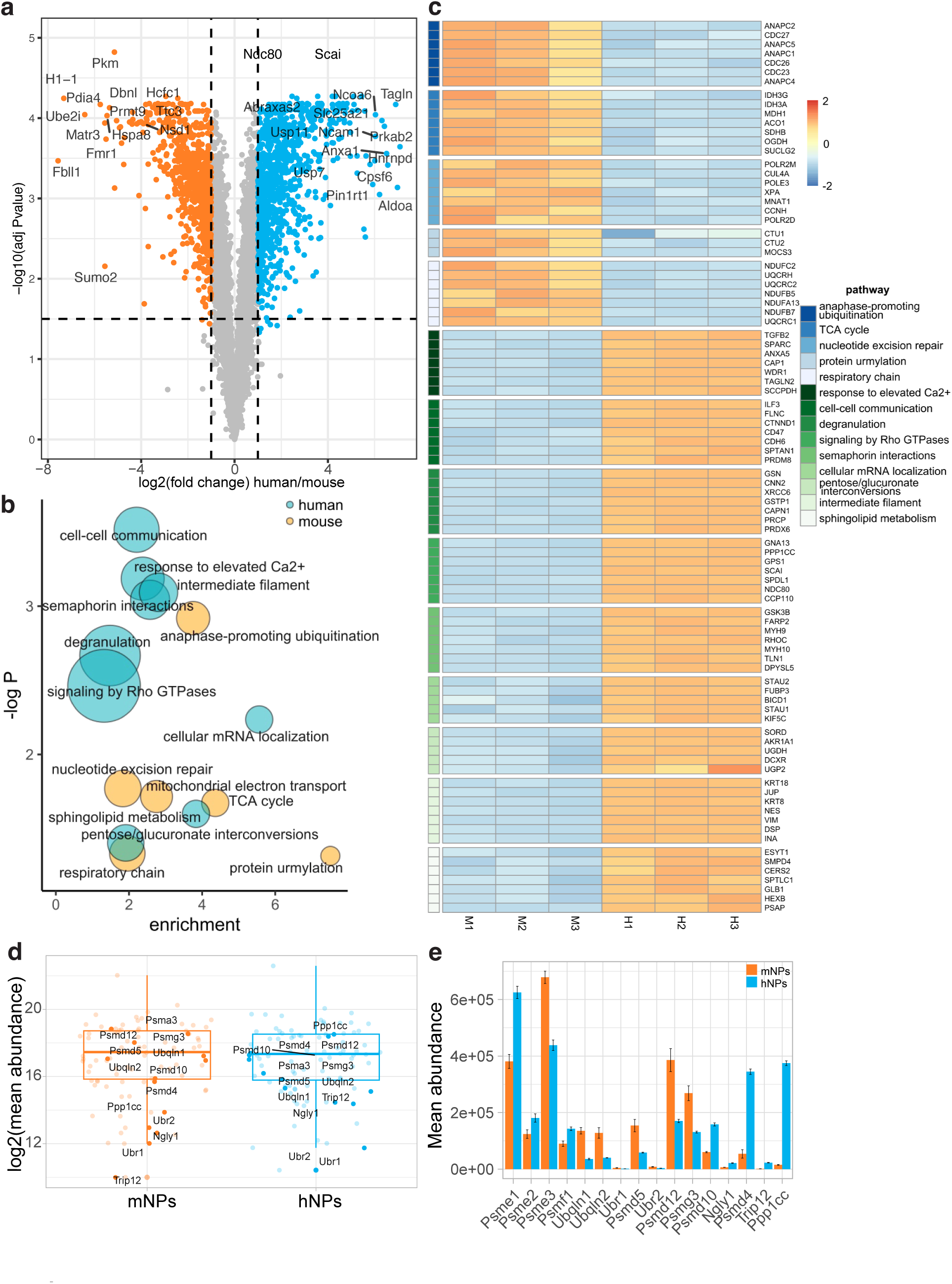
Global proteomic differences in mouse and human neural progenitors. (a) Volcano plots showing 1791 Differentially Expressed Proteins (DEPs) with ≥ 2 FC and FDR ≤ 0.05 in at least 2 biological replicates. DEPs enriched in mouse or human are depicted in orange (794) and blue (997), respectively. The full list of DEPs is presented in Table S2. (b) Bubble plot depicting the top pathways identified in pathway analysis with DAVID (Table S3). The pathways enriched amongst human- or mouse-upregulated DEPs are shown in blue and orange, respectively. Bubble sizes indicate the number of DEPs in each pathway. The axes show the fold enrichment of each term compared to the background, and - log of P value from a Fisher’s test. (c) Heatmap of the normalized scaled intensities of DEPs assigned to Gene Ontology categories (DAVID) in mouse and human NPs. Each pathway is color coded and ordered by p value. (d) Abundance of proteins associated to the ubiquitin-proteasome pathway in mouse and human NPs. DEPs are highlighted in darker color and labeled. (e) Mean abundance of DEPs associated with the ubiquitin-proteasome pathway in mouse and human NPs.

Many of the differentially expressed proteins were related to the ubiquitin–proteasome system and ubiquitination (Fig. 2d; Table S2), and many proteasomal subunits were more abundant in mouse (Fig S2g, h). There were species-specific differences in subunits of the PA28 proteasome regulatory particles, with human NPs having increased levels of PA28αβ (PSME1/E2), and mouse NPs containing higher PA28γ/Psme3 levels (Fig. 2e). Both PA28αβ and PA28γ particles have been shown to be less efficient than the 19S canonical regulatory particle at activating the 20S proteolytic activity^18^, and could compete with one another for binding to this 20S core. Proteasome-related proteins such as Ttc3, Psma3, Psmd5, Psmd12; ubiquitin ligases Smurf1, Rnf34 and Cul7; F box protein Tbl1x; proteasome accessory protein Ubqln1, Ubqln2, Ubr1, Psmd5, Ubr2, Psmd12, and Psmg3 and zinc finger protein Zfand1 were upregulated in mouse NPs, supporting a potential increase in protein degradation in mouse compared to human NPs. Proteins related to sumoylation (Ube2i, Sumo2), and enzyme transferases (Prmt9, Nsd, Hcfc1) were detected as differentially increased in mouse NPs as well (Fig. 2a). By contrast, the chaperones and regulatory subunits PSMD10, NGLY1, PSMD4, TRIP12, and PPP1CC were significantly upregulated in human NPs (Fig. 2c, d). The expression of deubiquitinases USP7, USP11, and ABRAXAS2, implicated in the stabilization of proteins^19,20^, were increased in human NPs.

To explore further the differences between mouse and human cells we performed pathway enrichment analysis^21^ (Table S3). The pathways enriched in mouse were the anaphase-promoting ubiquitination complex, a major E3 ubiquitin ligase complex with roles in the control of cell cycle and neurogenesis^22–24^, the mitochondrial electron transport chain, specifically complex III/ Ubiquinol-Cytochrome C Reductase, and enzymes of the TCA cycle (e.g. isocitrate dehydrogenase subunits IDH3G and IDH3A) (Fig. 2b,c). The pathways enriched in human NPs represented upregulation of cell-cell communication (e.g., NCAM1, log2FC=6; PODXL), Ca2+ signalling, immunity and secretion (e.g., GSN, SRP14), as well as Rho GTPase-mediated signal transduction (e.g., GNA13, GNAI1). Specific human-upregulated pathways included semaphorins, involved in the axonal development (e.g., DPYSL5 hydrolase and dihydropyrimidinase CRMP1), as well as sphingolipid metabolism (e.g., SMPD4), involved in the neuronal development^25,26^. The Pentose/Glucoronate interconversion pathway was upregulated in human cells. Localized in the cytoplasm, this pathway is a key route to carbohydrate synthesis. It also plays a central role in recycling NADP to NADPH, a cellular reducing agent for reductive biosynthesis and protection against oxidative stress^27^.

Together, the comparison of the mouse and human proteome in NPs indicated selective upregulation of proteins related to degradation and mitochondrial metabolism in the mouse. Proteins overrepresented in human were associated with neural phenotypes, the biosynthesis of carbohydrates, and the stabilization of the proteome.

### Systematic differences in the fold-change stability of protein orthologs

Having established a genome-wide difference in protein turnover between mouse and human NPs^4^, we proceeded to profile protein turnover in mouse and human to determine the identity of specific proteins and pathways that exhibit similar or divergent protein dynamics between species. We performed dynamic SILAC (Stable Isotope Labeling by Amino acids in Cell culture) coupled to mass spectrometry in mouse and human NPs to estimate half-lives of individual proteins at a proteome-wide scale^28^. The labelling was centered on equivalent time points in mouse and human NPs based on our previous stability analysis^4^, and the collection of five time points per species was distributed according to the expected stability differences between mouse and human. The media was switched to heavy lysine and arginine isotopes 24 hours before day 2 in mouse. For human, we started labeling on day 4, 48 hours before human NPs reached the stage assayed (Fig. 3a). The samples were analyzed by LC-MS/MS at each time-point, and the degradation rate constants for individual proteins calculated (see Methods). We determined the half-lives of 4990 and 4640 proteins in mouse and human, respectively (Fig. 3b, Table S5, S6). Their median half-lives were 18.2 hours and 27.9 hours, respectively. To obtain more detailed insight into protein features associated with differences in stability in the mouse and human datasets, we performed gene set enrichment analysis (GSEA) across all GO terms (Fig.S3a,b). Structural proteins were enriched for long half-lives in both mouse and human, as expected. Furthermore, proteins with metabolism- and translation-related functions, and proteins located in the mitochondrial and ribosomes were more stable on average, especially in mouse (Table S7, S8).

**Figure 3.**
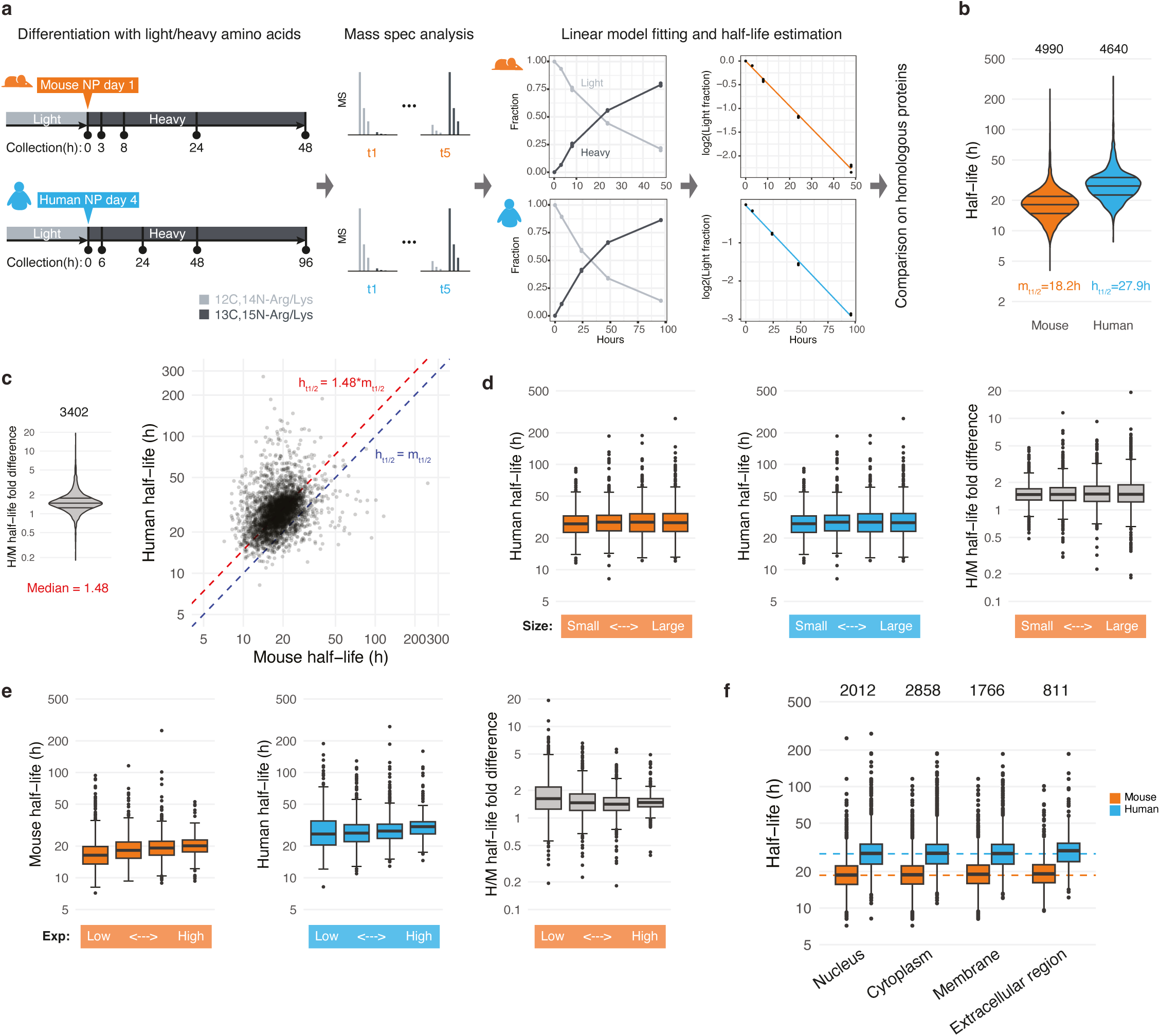
Dynamic SILAC proteomics identifies global differences in protein stability. (a) Schematic of dynamic SILAC combined with quantitative proteomics. (b) Violin plot showing the estimated half-lives of mouse and human proteins in NPs. Median half-lives for the mouse and human datasets are indicated. Quartile lines in the violin plots are the 25th, 50th (median) and 75th percentiles.= (c) Violin plot illustrating the fold change in half-lives between human and mouse proteins (left), and dot plot showing the half-lives of 3402 homologous proteins in mouse and human (right). The blue line represents a 1:1 relationship, indicating identical half-lives in both species. The red line marks the median fold-difference in half-life between human and mouse proteins. Violin plot as in (b). (d) Half-life measurements of homologous proteins divided into quartiles based on protein size (amino acid sequence length), from smallest to largest. Orange and blue box plots represent the values for mouse and human proteins, respectively. Grey box plots indicate the fold difference in half-lives in human versus mouse. Box-plot: Centre line is the median; box represents upper and lower quartiles; whiskers are1.5x interquartile range from the upper and lower quartiles; dots are outliers. (e) Quartiles of measured half-lives homologous proteins partitioned according to protein abundance (average total expression level), from low to high. Orange and blue box plots correspond to mouse and human values selected for partitioning, respectively. Grey box plots indicate the fold difference in half-lives in human versus mouse. Box-plot as (d). (f) Half-lives of homologous proteins associated with four GO terms in cellular component. Dashed lines indicate median half-lives of mouse and human homologues. Box-plot as in (d).

We further compared the half-lives of 3402 homologous proteins detected in both mouse and human datasets (Fig.3c, Table S9). Half-lives of human homologues were generally longer than those of mouse homologues, showing a median fold difference of 1.48 (Fig. 3c), consistent with the measurement for OLIG2 (Fig. 1a). To investigate whether specific features of proteins determine their stabilities, we first split the 3402 homologous proteins into quartiles based on their size (amino acid sequence length) or abundance (average total expression level). Each quartile exhibited similar species-specific trends of half-lives and interspecies fold differences despite their sizes (Fig. 3d; Fig. S3c,d). We observed a mild correlation between protein abundance and half-lives, with more abundant proteins tending to be more stable; however, the half-life fold differences between species were similar irrespective of expression levels (Fig. 3e; Fig. S3e,f). These results indicated that the overall differences in protein stability between mouse and human NPs are independent of protein size and abundance.

To examine if differences in the subcellular localization of proteins affected half-life, we analysed the dataset categorised by cellular compartments (see Methods). Proteins assigned to the nucleus, cytoplasm, membrane or extracellular compartment showed similar median half-lives in each species (Fig. 3f). Consequently, the observed stabilisation of the human proteome compared to the mouse occurs in all cellular compartments. We then categorised proteins based on their molecular functions by referring to GO terms (Fig. S3g). Although most of the groups followed the overall trend, proteins classified as “Structural molecule activity”, such as TUBB, RPL32 and LMNA, and “ATP-dependent activity” appeared longer-lived, while the groups “Transcription regulator activity”, including transcription factors like POU3F2, YAP1 and ZEB2, and “Molecular adaptor activity” were short-lived in comparison with the overall median (Table S10). Nevertheless, the fold difference in half-life between the species was preserved for the proteins. Together, the results indicate that proteins in diverse biophysical properties, functional categories, and subcellular locations have similar stability differences between mouse and human NPs.

Given that our earlier comparative proteomics analysis had indicated several differences in the proteasome between the two species, we reasoned that the observed differences in proteome half-lives could result from differences in the stability of specific proteasome components, which together accounts for ∼70% of the cellular protein degradation^29^. We grouped proteins associated with the proteasome into three categories: core proteasomal components, proteasomal regulatory components, or proteasomal activators/inhibitors. The classification was based on a curated list generated by the proteostasis consortium^30^. We then compared the median half-lives of these protein categories with the median proteome half-life in mouse and human.

Core particles followed the median trend for mouse and human (Fig. S3h). By contrast, regulatory particles were more stable than average in both species, especially in mice (Fig. S3h). For proteins in the “activator/inhibitor” category, human proteins were less stable relative to the human median, but this pattern was not observed in mouse (Fig. S3h, i). For example, the half-life of human PSME3 was shorter than the human median, whereas mouse Psme3 aligned with the mouse median (Fig. S3i). The increased degradation rate in human PSME3 explains the significant enrichment of Psme3 in mouse in the global dataset (Fig. 2e).

Similarly, the proteasome regulator Psmf1^31^ was less stable than the mouse median, but human PSMF1 showed stability comparable to the human median. Global proteomic quantification indicated that Psmf1 was expressed at low levels overall, but at higher levels in human than in mouse (Fig. 2e), consistent with the measured dynamics. Together, these results identify species-specific differences in the stability of proteasome activators and inhibitors. They suggest that the differential availability of these regulators could contribute to the differences in protein stability between mouse and human.

### A role for active protein degradation in mouse and human neural progenitors

The faster cell cycle in mouse compared to human^4^ raised the possibility that the lower rate of protein turnover in human NPs could arise from lower cell division rates resulting in a lower rate of passive dilution. To test this, we set out to compare protein stability in non-dividing cells by inducing the generation of post-mitotic neurons *in vitro* from mouse and human NPs (Fig. 4a,b). Treatment with the gamma-secretase inhibitor Dibenzazepine (DBZ), which inhibits the activation of Notch pathway, increased the production of TUBB3+ neurons at the expense of SOX2 expressing NPs^32^ (Fig. 4a, Fig. S4a, b). Measurements of protein stability in these conditions by AHA pulse-chase labeling^4^ showed that the half-life in mouse post-mitotic neurons was ∼10h (t1/2 = 11.45 ± 1.8 hours), and ∼20h in human neurons (t1/2 = 22.93 ± 4.58 hours) (Fig. 4c,d, Fig. S4c). These rates were similar to those measured in NPs (mouse NPs t1/2 = 9.3h ± 1.99 hours; human NPs t1/2 = 17.57h ± 3.6 hours). These results indicate that protein degradation is slower in human neurons than mouse neurons and support the conclusion that active protein degradation mechanisms drive differences in protein stability between mouse and human.

**Figure 4.**
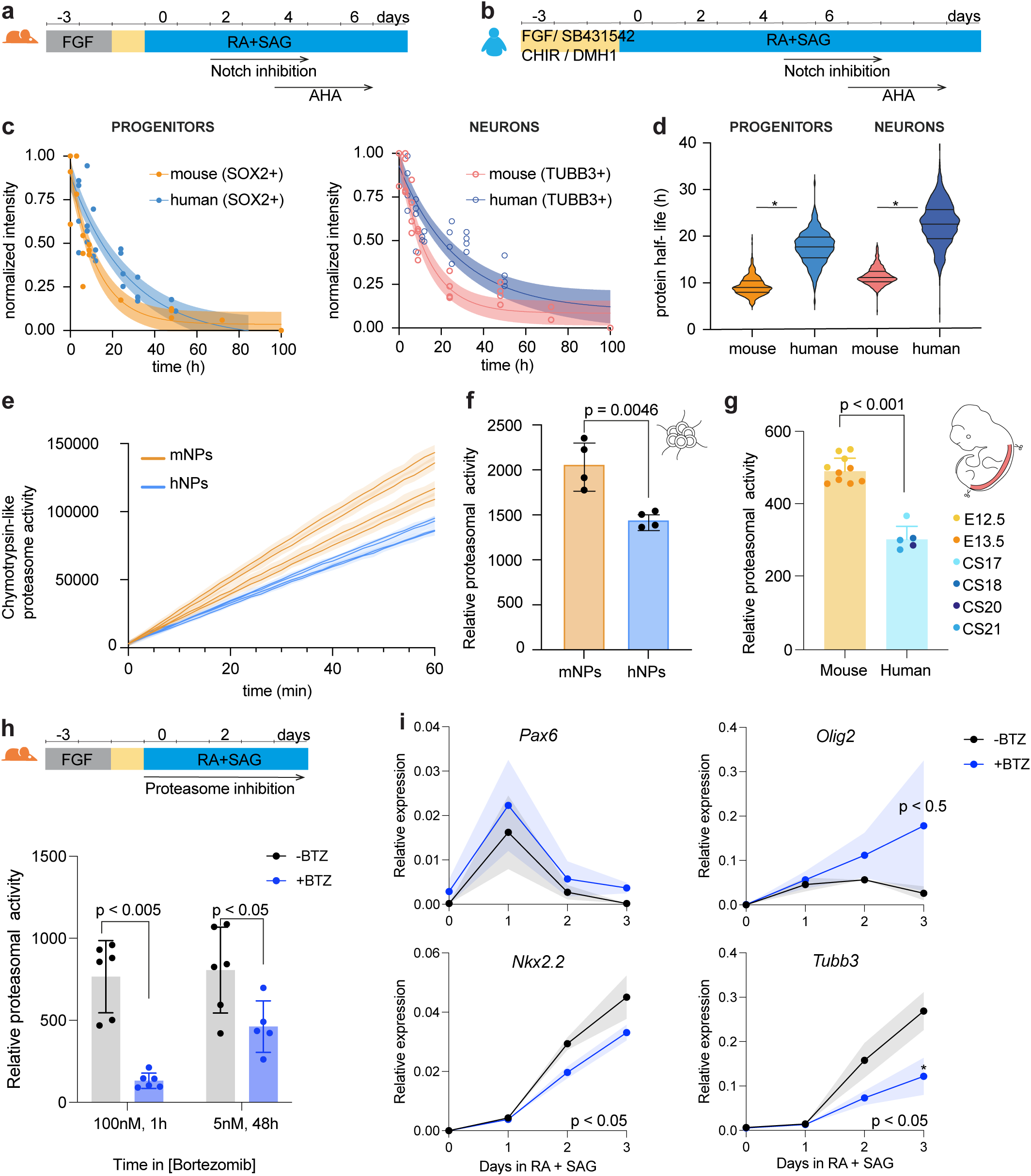
Proteasomal activity is higher in mouse neural progenitors. (a, b) Schema of the experimental design for neuronal differentiation from ESCs. A Notch inhibitor (10mM DBZ, gamma secretase inhibitor) was added to increase the differentiation of NPs into post-mitotic neurons. In mouse, treatment was administered from day 2 to day 4 of the differentiation. On day 4, AHA pulse and chase experiments were performed to measure protein stability in progenitors and differentiated neurons in mouse (a). Similarly, human NPs were treated for 48 hours with the Notch inhibitor and protein stability measured after treatment starting at day 4 (b). (c) Normalized intensity of AHA levels in mouse and human NPs (mouse, orange; human, blue) and post-mitotic neurons (mouse, red; human, dark blue) across timepoints. Curves and shadowed areas show best exponential fit with 95% confidence intervals; (mouse n = 2, human n = 2 in duplicates). (d) Half-live estimation of the proteome in mouse and human neural progenitors (mouse, orange; human, blue) and post-mitotic neurons (mouse, red; human, dark blue) for datasets in (c). Quartile lines in the violin plots are the 25th, 50th (median) and 75th percentiles. Statistical significance (*) corresponds with < 5% of overlap between the distributions of parameter estimations (mouse n = 2, human n = 2 in duplicates). (e) Fluorescence intensity measurements of the chymotrypsin-like enzymatic activity of the proteasome using a fluorescent peptide substrate. A line connects the mean values at the measured timepoints, and the shaded areas represent the standard deviation (n=4). (f) Proteasomal activity measured as the slope of fluorescence increase from substrate digestion in (e) in mouse and human NPs. Data are presented as mean ± standard deviation; p-value is from unpaired t-test (n=4). (g) Slopes of fluorescence increase from the digested substrate as the proxy of proteasomal activity in spinal cords of mouse and human embryos. Mouse embryonic day (E) 12.5 and 13.5; Human stages span Carnegie Stages (CS) 17-21. Data are presented as mean ± standard deviation; p-value is from unpaired two-tailed Welch t-test (mouse n = 10. E12.5 (4) and E13.5 (6); human n = 5. CS17 = 1, CS18 = 1, CS20 =2, CS21=1). (h) Schema of the experimental design for Bortezomib (BTZ) treatment on mouse neural differentiations from ESCs, and measurements of proteasomal activity at the indicated timings and concentrations. Data are presented as mean ± standard deviation; p-value is from paired t-test (n=3 in duplicates). (i) RT-qPCR analysis of *Pax6*, *Olig2*, *Nkx2.2* and *Tubb3* expression in mouse neural differentiations treated with Bortezomib (+BTZ) compared to controls (-BTZ). Data represent mean ± standard error of the mean; two-way ANOVA followed by with Sidak’s multiple comparisons test. Reported P-values correspond to the ANOVA results, while asterisks indicate statistical significance from the post hoc test (n = 4).

A role for active protein degradation in the species differences in protein turnover predicts higher proteasome activity in mouse NPs than human NPs. This is supported by the global and SILAC proteomic analyses that indicated differences associated with protein degradation (Fig.2b, Fig. S3h, i). To test this directly, we used a fluorogenic assay to measure chymotrypsin-like proteasomal activity^33,34^ in mouse and human NPs at equivalent developmental stages, normalized to total protein amount (Fig. 4e). We observed 1.43-fold higher activity in the mouse compared to human (Fig. 4f). A similar difference was also observed in two different mouse and human embryonic stem cell lines (Fig. S4d). Moreover, assays of proteasomal activity from *in vivo* samples of mouse and human developing spinal cords indicate higher levels of proteasomal activity in mouse compared to human (Fig. 4g, S4e). This is consistent with the idea that higher proteasomal activity in mouse spinal cord progenitors compared to human contributes to the higher rate of protein degradation in mouse.

To test if decreasing proteasomal activity leads to a delayed differentiation in mouse, we treated mouse differentiations with low doses of the proteasome inhibitor Bortezomib (BTZ) from the time of RA and SAG addition. Cells treated for 1 hour with 100nM of BTZ showed a significant decrease of proteasomal activity but died within 24h. Cells treated with 5nM BTZ survived for four days and showed decreased proteasomal activity, albeit less marked than for 100nM BTZ (Fig. 4h). Nevertheless, there was a decrease in the percentage of NPs (SOX2+) and neurons (TUBB3+) 72 - 96 hours after treatment suggesting deleterious effects of prolonged treatment (Fig. S4f,g). This is possibly due to proteotoxic stress, and limited our analysis on the pace of differentiation. The fraction of early neural progenitor markers PAX6+ and OLIG2+ cells was unchanged upon 5nM BTZ treatment (Fig. S4h,i), but RNA levels and protein intensities were higher for the two markers up to 72 hours after treatment in the few surviving cells (Fig. 4i, S4j,k). In contrast, the expression of the late differentiation markers *Tubb3* and *Nkx2*.2 were decreased (Fig. 4i). This indicates a moderate stabilization of early transcription factors within the gene regulatory network in response to reduced protein degradation via proteasomal inhibition, accompanied by a slowdown of mouse motor neuron differentiation.

### IRX3 depletion accelerates OLIG2 expression

Although global proteasome inhibition suggested that an increased stabilization of the proteome decelerates mouse motor neuron differentiation, the associated proteotoxic stress limited the interpretation. Thus, we sought to test whether decreasing the stability of a specific regulatory protein accelerated the tempo in NPs. We focused on IRX3, which is expressed early in spinal cord development, prior to cells receiving sonic hedgehog signaling, and represses OLIG2 and motor neuron progenitor formation^9,35^. To this end we generated an Irx3::HA::HALO mouse ESC line (Irx3-HALO) by inserting the HALO tag at the C-terminus of the Irx3 gene. An HA epitope was inserted as a linker between the Irx3 coding region and HaloTag^36^.

We tested Irx3-HALO expression and functionality in spinal cord motor neuron differentiation. We differentiated Irx3-HALO cells, and we exposed cells to RA only on day 3 to promote IRX3 expression (Fig. S5c). We then performed a pulse-chase experiment with HALO-JF549 ligand (Fig. S5a). Cells were pulsed for 1h, ligand was then removed, and samples collected 0h, 4h and 24h later. We observed decreasing levels of HALO-JF549 signal during the chase, while the level of IRX3 expression, assayed by HA immunofluorescence, remained constant across the time course (Fig. S5b, c). This confirmed that the Irx3-HALO line reproduces the expected expression pattern of IRX3 and validates the use of the HA::HALO tag as a multimodal tag to track IRX3 levels and dynamics of turnover.

The HALO tag allowed us to use a small molecule degrader, HaloPROTAC, to induce IRX3 degradation. HaloPROTAC binds covalently to the Halo tagged protein and recruits specific E3 ligases, which promote ubiquitination of the protein and subsequent degradation by the 26S proteasome^36,37^ (Fig. 5a). To test whether HaloPROTAC decreased IRX3 protein stability, IRX3 expression was induced by addition of RA for 24h as before, and cultured thereafter in medium with RA and SAG. HaloPROTAC was added to the Irx3-HALO at the time of RA addition. We collected samples at 0h, 12h, and 24h after HaloPROTAC treatment as well as in a control Irx3-/-cell line. While IRX3 was detected across time points in the untreated control condition (-PROTAC), IRX3 was reduced 24h after HaloPROTAC addition (0h time point), and HA intensity was reduced almost to the levels found in the Irx3-/-line 48h after HaloPROTAC addition (Fig.S5d). These results confirm protein depletion of by addition of HaloPROTAC.

**Figure 5.**
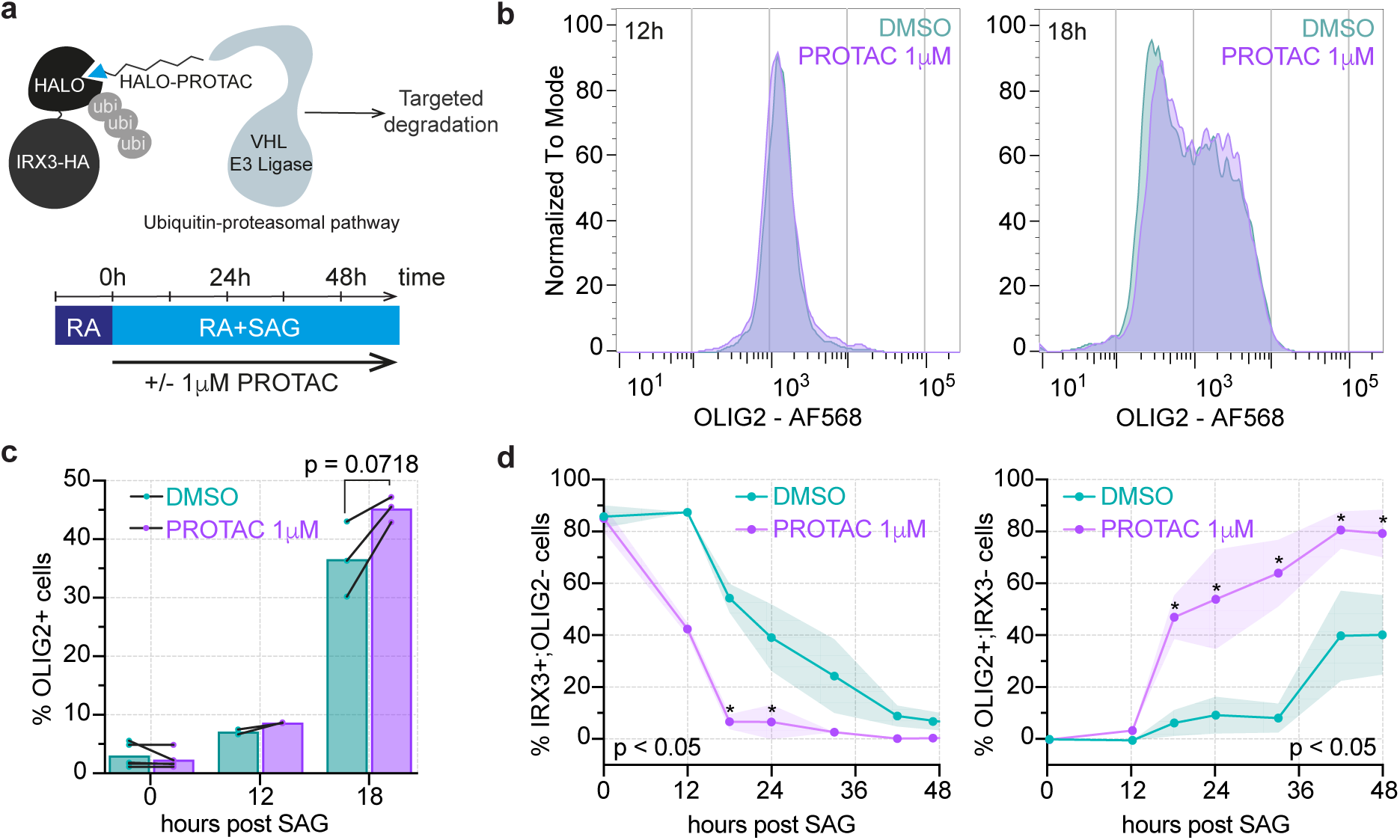
HALO-PROTAC addition depletes IRX3 and increases the proportion of OLIG2 expressing neural progenitors. (a) Schematics of HaloPROTAC targeted degradation and treatment regimes. HaloPROTAC covalently binds to HALO and induces the 26S proteasomal degradation of IRX3-HALO through the recruitment of the von Hippel-Lindau (VHL) E3 ubiquitin ligase. (b) Representative FACS plots of OLIG2 expression measured in SOX2 expressing mouse NPs in controls (DMSO, turquoise) and PROTAC (+PROTAC, violet) treated samples at 12h and 18h post addition. (c) Percentage of total OLIG2 positive mouse NPs at 0h, 12h and 18h post SAG addition in controls and PROTAC treated samples. Data lines represent biological replicates; p-value is from paired sample t-test (0h n=5; 12h n=2; 18h n=3). (d) Percentage of IRX3+;OLIG2- and OLIG2+;IRX3-NPs during differentiation in DMSO or with PROTAC samples. A line connects the mean values (dots), and the shaded areas represent the standard deviation. Two-way ANOVA followed by with Sidak’s multiple comparisons test. Reported P-values correspond to the ANOVA results, while asterisks indicate statistical significance from the post hoc test (n =5 except 12h n=2 and 18h n=3).

We next tested how acute IRX3 degradation impacted the dynamics of OLIG2 expression, a target of IRX3 repression^10,38^. We cultured differentiating Irx3-HALO cells for 24h in RA to induce *Irx3* in neural progenitors as above. Cells were then switched to media containing RA and SAG either in the presence or absence of HaloPROTAC degrader. This culture regime allowed the maximal expression of IRX3 prior to OLIG2 expression induced by SAG induction. We assayed OLIG2 expression 12-48h after SAG and HaloPROTAC addition (Fig. 5b, c). In NPs that had not been exposed to HaloPROTAC, SAG-dependent repression of IRX3 and induction of OLIG2 took more than 18h. By contrast, in cells treated with HaloPROTAC, IRX3 was largely absent by 18h and an increase in the percentage of OLIG2+ cells was apparent (Fig. 5d). Quantifications of the IRX3-;OLIG2+ population revealed a higher proportion of OLIG2+ cells and lower variability in the HaloPROTAC condition compared to control (Fig. 5d). Moreover, fewer cells co-expressed IRX3 and OLIG2 with HaloPROTAC, indicating a more rapid transition from the IRX3+;OLIG2-to an IRX3-;OLIG2+ state (Fig. S5e). Together, these results demonstrate that acute degradation of the transcriptional repressor IRX3 accelerates the appearance of OLIG2 expressing neural progenitor cells, providing evidence that increasing protein degradation can accelerate the tempo of developmental gene regulatory networks.

## DISCUSSION

Here we provide a systematic comparison of protein dynamics between mouse and human neural progenitors. Through a combination of targeted protein labelling, quantitative mass spectrometry, and genetic manipulation of endogenous proteins, we have characterised protein production and degradation rates in equivalent mouse and human NPs. The data provide insight into the mechanisms underlying the differences in developmental tempo between these species.

Our findings show a ∼1.5 fold-difference in the stability of homologous proteins between mouse and human NPs. The trends we observe are consistent with previous assays that measured the overall degradation rate of the proteome by global metabolic labeling^39–43^. Alongside the differences in protein degradation rates between species, we find a ∼2 fold-difference in the rate of production between mouse and human NPs. The difference in protein production rates between species suggests that overall protein metabolism (production and turnover) is a primary driver of the observed tempo differences. Protein synthesis and degradation rates are key determinants of overall protein concentration and shape tissue-specific proteomes in adult cells^43–45^. Yet, their roles in developmental systems, where cells simultaneously change size and growth rate, remain poorly understood. Differences in protein translation and degradation rates between species result in overall similar protein concentrations in human and mouse NPs as human cells are of bigger size. Comparable protein densities in the two species have also been reported for presomitic cells^5,6^. These observations suggest that differentially expressed proteins involved in basal biochemical kinetics may underlie the differences in developmental tempo between mouse and human.

Our findings further support a role for active protein degradation in controlling developmental tempo. Unbiased global proteomic analysis suggests an increased abundance of proteins associated with degradative pathways and metabolic functions. Importantly, we found that the differences in protein stability persist in post-mitotic neurons, indicating that active degradation, rather than passive dilution through cell division, is a primary mechanism driving the differences in species-specific protein turnover rates. The increased stability of human proteins in NPs was observed across most cellular compartments and was independent of amino acid sequence, protein size or abundance. At least part of this differential protein degradation rates between mouse and human NPs can be explained by differences in proteasomal activity. We observed ∼1.5-fold higher chymotrypsin-like proteasomal activity in mouse NPs compared to human NPs. Importantly, we also detected ∼1.73-fold higher activity in mouse embryonic spinal cord tissue relative to stage-matched human *in vivo* samples, indicating the species difference in proteasome activity is also found *in vivo*.

The species-specific differences in proteasomal activity may be regulated either by differences in the abundance or activity of regulatory complexes and/or proteasome particles, through the availability of proteasome targets, or through other post-translational regulatory mechanisms. In line with this, our comparative proteomics identified a selective upregulation of proteins related to degradation, with differences between the two species in ubiquitin-proteasome pathway components that could have roles in each of these facets of proteasomal regulation. For example, we identify differences in the abundance and stability of regulatory particles of the PA28αβ and γ complexes. While the PA28αβ is thought to be characteristic of the immune system, PA28γ is expressed in all cell types. Also, PA28 complexes promote protein degradation in an ATP and ubiquitin-independent manner^18,46–48^. While there did not appear to be differences in the abundance of all core particle proteasome subunits between species, quantifications of proteasomal abundance in their native configurations as well as measurements of assembly rates will provide a better mechanistic interpretation for the measured differences in proteasomal activity. In the future, dissecting the functional outcomes of the different proteasomal regulatory complexes in mouse and human NPs will be key to understand how homologous proteins are degraded at different rates.

The importance of protein degradation in regulating developmental pace was tested by the treatment of mouse NPs with low doses of a proteasome inhibitor. Whilst the differentiation was strongly affected by the treatment, early motor neuron progenitor markers displayed reduced intensity at early time points and persisted modestly beyond their expected temporal window in surviving cells, whereas late differentiation markers were expressed at lower levels. These findings suggest that decreased proteasomal activity slows developmental tempo. However, because the proteotoxic effects of proteasome inhibition limited the interpretation of these results, we also examined the effect of protein degradation by the targeted manipulation of IRX3, a key transcriptional repressor in neural progenitor pattern. Increasing the degradation rate of IRX3 using a HaloPROTAC system led to accelerated expression of its target gene, OLIG2, and suggests an accelerated transition of intermediate NPs to ventral motor neuron progenitors. This demonstrates that modulating the stability of individual regulatory proteins can have a significant effect on gene expression dynamics and cell fate transitions during development. Moreover, these experiments offer proof-of-concept for the use of the HaloPROTAC system for targeted perturbation of components of a gene regulatory network that could be broadly applied to dissect gene regulatory mechanisms.

Consistent with our findings, Matsuda et al. report a pervasive ∼1.5-fold slower rate of protein degradation in presomitic mesoderm^49^, and Swovick et al. describe differences in protein turnover across twelve different mammalian species with divergent lifespan^50^. The conservation in different tissues of species-specific differences in protein degradation rates suggests that this may be a fundamental mechanism for evolutionary adaptation of timing in development and homeostasis. Our work measures differences in abundance and stability of regulatory proteasomal subunits that may have a global impact on tempo. Future studies investigating the molecular basis for the increased protein production and proteasomal activity in mouse cells, as well as potential differences in other degradation pathways such as the autophagy-lysosome system, will be important for fully understanding these species-specific differences.

While much attention for the basis of interspecies differences has focused on the regulatory logic of gene networks through gene duplication or de novo acquisition of enhancers^2,51,52^, our results emphasise that post-translational processes, particularly protein metabolism, play a crucial role in shaping the temporal dynamics of development. Modulating protein stability could offer a flexible and precise method for globally controlling developmental tempo, without requiring extensive changes to gene regulatory networks or protein sequences in individual developmental processes. Whether differences in protein production and degradation rates also account for heterochronic changes in developmental pace, such as those associated with the expansion of the primate cortex or the protracted maturation of cortical neurons remains to be determined^3,52,53^. Our work highlights the importance of considering post-translational regulation when studying developmental systems and evolutionary divergence between species. Little is known about the mechanisms that drive and maintain the homeostasis of proteins (proteostasis) across species and how they affect gene regulatory programs in development. Exploring proteostasis in different evolutionary contexts could provide insight into the constraints and selective pressures shaping developmental timing across species.

In conclusion, our study provides a comparison of protein dynamics between mouse and human NPs, revealing that differential protein degradation appears to be a key driver of species-specific developmental timing. These findings raise questions about the molecular mechanisms underlying evolutionary changes in developmental tempo and the potential role of proteostasis in other aspects of phenotypic diversity between species. Future studies exploring how protein production and degradation are controlled globally and its impact across developmental processes in diverse organisms will be essential for deepening our understanding of the molecular basis of evolutionary change. Understanding these mechanisms will not only contribute to our knowledge of evolutionary and developmental biology but may also inform strategies for manipulating cellular timing in contexts such as stem cell differentiation, regenerative medicine, and homeostasis.

## MATERIALS AND METHODS

### Cell culture and neural progenitor differentiation

The use of human ESCs was carried out in accordance with approvals from the UK Stem Cell Bank Steering Committee (SCSC14-18 to J.B. and SCSC21-46 to T.R.). All cell lines used in this study were confirmed to be mycoplasma negative. WA09/H9 ESC line (WiCell), and OLIG2::HA::SNAP H9 ESC lines were routinely cultured in Essential 8 medium (Thermo Fisher A1517001) or StemFlex medium (Thermo Fisher A3349401) on 0.5 µg/cm^2^ laminin-coated plates (Thermo Fisher A29249) or 0.5 µg/cm^2^ of Vitronectin (Thermo Fisher A14700), and split using Versene (Gibco 15040066).

Mouse ESCs (HM1, OLIG2::HA::SNAP and IRX3::HA::HALO mouse ESC line) were propagated on mitotically inactivated mouse embryonic fibroblasts (feeders) in DMEM knockout medium supplemented with 1000U/ml ESGRO-LIF (ESG117 Sigma Aldrich), 10% cell-culture validated fetal bovine serum, penicillin/streptomycin, 2 mM L-glutamine (GIBCO) or on 0.1% gelatine in ES medium supplemented with LIF/2i, consisting of 1 μM PD0325901 and 3 μM CHIR99021P (Stem Cell Institute).

Mouse and human NP differentiation was performed following Rayon et al.^4^. For cells grown on ES medium with LIF/2i, 2i was removed at the split prior to differentiation. The γ-secretase inhibitor DBZ was used in experiments to fully differentiate neurons. Cells were treated with 10 ng/uL DBZ (Tocris Biosciences Cat. No. 4489) for 72h starting at day 2 for mouse and day 5 in human. The proteasome inhibitor bortezomib (R&D, 7282) was used in experiments to stabilize the proteome starting at day 0 after RA and SAG addition at the indicated concentrations. Samples were collected at the indicated time points and processed for intracellular flow cytometry (see below). Medium was changed daily.

For IRX3::HA:: HALO mESC differentiation, N2B27 was supplemented with 100 nM RA (R2625 Sigma Aldrich) for neural induction at day 0 and supplemented with RA and 500 nM SAG (566660 Calbiochem) thereafter. To achieve IRX3 depletion, from day 1 onwards, cells were treated with 1 mM HaloPROTAC3 (Promega GA3110) or DMSO as control. Samples were collected after 12h, 18h, 24h, 33h, 42h, 48h of HaloPROTAC or DMSO treatment and processed for intracellular flow cytometry (see below). Medium was changed daily.

### Generation of mouse and human ESC lines by CRISPR

All transgenic cell lines were generated by CRISPR-Cas9–mediated homologous recombination, as in Gouti et al. 2017^54^ and Rayon et al. 2020^4^. Briefly, 2 × 10^6^ cells were electroporated with 2 μg of each plasmid using program A23 of Nucleofector II (Amaxa) and mouse stem cell Nucleofector kit (Lonza DPH-1001) or human stem cell Nucleofector I kit (Lonza VPH-5012). For selection, colonies were first treated with 0.5 mg/ml Puromycin (Sigma P9620) for 2 days followed by 50 μg/ml Geneticin (Thermo Fisher 10131027) selection. Individual colonies were picked and replated to allow expansion and a second round of Geneticin selection. Correct integration of the T2A-HA-SNAPTag transgene was verified using long-range PCRs and Sanger sequencing.

#### Generation of the pNTKV-HA-HaloTag and pNTKV-HA-SnapTag vectors

The coding sequence for the HaloTag was extracted by 2-step PCR amplification with Phusion High-Fidelity DNA Polymerase (NEB, M0530) from the pFC27A HaloTag CMV-neo Flexi vector (Promega, G842A), while, at the same time, the coding sequence for the HA epitope (tatccctatgacgtcccggactatgca) and two stop codons were introduced as part of the 5’ primer used for the amplification process.

For the HA-SnapTag cell line, the coding sequence for the SnapTag was extracted equally by PCR amplification from the pSNAPf commercialvector (NEB, N9183S), while the HA coding sequence and two stop codons were introduced as part of the 5’ primer.

The resulting purified DNA elements were separately cloned, by restriction digestion and inserted with in-Fusion cloning into the pNTKV-T2A-mKate vector^55^.

#### Generation of Olig2::HA::SNAP mouse and human ESC lines by CRISPR

For CRISPR-Cas9–mediated homologous recombination, short guide RNA (sgRNA) sequences (mouse: CGGCCAGCGGGGGTGCGTCC, human: CTGTCGCCAGAACGCGC) were cloned into pSpCas9(BB)-2A-Puro (Addgene pX459 plasmid no. 62988). As donor vector, the HA-SnapTag cassette was inserted at the 3′ end of the Olig2 open reading frame, using 2.83-Kb upstream and 5.04-Kb downstream arms for mouse and 2.78-Kb upstream and 1.98-Kb downstream homology arms for human.

#### Generation of mouse Irx3::HA::HALO ESC line

For CRISPR-Cas9–mediated homologous recombination, short guide RNA (sgRNA) sequence: ATGGTTGAAAAGTTAAGACG was cloned into pSpCas9(BB)-2A-Puro (Addgene pX459 plasmid no. 62988). As donor vector, the HA-HaloTag cassette was inserted at the 3′ end of the Irx3 open reading frame, using 2.32-Kb upstream and 0.34-Kb downstream homology arms.

#### Generation of mouse Irx3 knockout line

For CRISPR-Cas9–mediated non-homologous end joining to generate loss of function by frameshift two short guide RNA (sgRNA) sequences: TCTCTACCCACCCGAACGCC and GGATGTACTGGTATCCGAGC were cloned into pSpCas9(BB)-2A-Puro (Addgene pX459 plasmid no. 62988).

### SNAP/HALO labelling for pulse or pulse-chase experiments

For OLIG2:HA::SNAP experiments, mouse and human ESCs were differentiated to NP day 2 and day 6, respectively, as described above. To estimate OLIG2 degradation rates, OLIG2::HA::SNAP cells were incubated for one hour in RA and SAG medium containing 100 nM SNAP-Cell 647-SiR ligand (NEB S9102S). After two PBS washes, cells were incubated in RA and SAG medium containing 1µM SNAP-Cell Block (NEB S9106S) or 100nM SNAP-Cell TMR-Star ligand (NEB S9105S) for the indicated chase periods. To estimate OLIG2 production rates, cells were incubated for one hour in RA and SAG medium containing SNAP-Cell Block ligand. After two PBS washes, cells were incubated in RA and SAG medium containing SNAP 647-SiR ligand for the indicated chase periods.

For IRX3::HA::SNAP mouse pulse-chase experiments, cells were labeled for 1h in medium supplemented with 100nM Janelia Fluor HaloTag Ligand (549, Promega GA111) at day 0 on RA as described above. After two PBS washes, cells were incubated in RA and SAG medium for the indicated chase periods.

After SNAP/HALO ligand treatments, cells were washed once or twice with PBS for degradation or production measurements, respectively, and processed for intracellular flow cytometry or imaging. At least 3 biological replicates per species per time point from independent experiments were used. For OLIG2 protein dynamics, geometrical mean was measured on OLIG2 positive and negative populations. Half-life estimation was based on the non-linear model fitting in GraphPad Prism version 10.0.0 for MacOS using values normalized by samples not treated with SNAP-cell 647-SiR ligand. Error intervals reported correspond with 95% confidence intervals. Images were analysed in FIJI.

### Intracellular Flow Cytometry

Cells were dissociated with 0.5ml accutase (GIBCO) and fixed in 4% paraformaldehyde in PBS for 5min. For stainings, 0.5 - 1x10^6^ cells were used. Cells were incubated overnight with an antibody mix on PBST with 1% BSA at 4°C. The following day, and when HA (Cell Signaling C29F4, 1:1000) or OLIG2 (R&D AF2418, 1:800) primary antibody were used, cells were pelleted and incubated with donkey anti goat/rabbit Alexa Fluor secondary antibodies (1:1000) at room temperature for 1h. Cells were resuspended in 0.5 mL PBS and filtered for data acquisition on LSR Fortessa or Biorad ZE5 and MacsQuant Analysers. For OLIG2 measurements, 10,000 ∼ 50,000 total events were recorded. For other experiments 10,000 events gated on SOX2 were recorded. Analysis was performed using FlowJo software. The following fluorophore-conjugated antibodies were used: SOX2-V450 (BD 561610, 1:100), TUBB3-AF488 (BioLegend 801203, 1:100), TUBB3-AF647 (BioLegend 560394, 1:100), PAX6-647 (BD Biosciences 562249, 1:100).

### Global protein production measurements

#### AHA uptake experiment

Experiments were performed on differentiated neural progenitors (mouse day 2 and human day 6), as described above. The experiments were performed in the presence of methionine as we reasoned that methionine deprivation might trigger a reduction in protein production and impact protein translation. Spent medium was replaced by 1 mM AHA in differentiation medium supplemented with RA and SAG at 0h timepoint. Samples were collected at the indicated timepoints for intracellular staining with Click It Cell Reaction buffer kit (Invitrogen C10269) and Alkyne Alexa Fluor 488 (Invitrogen A10267) followed by SOX2-V450.

#### Puromycin uptake experiment

We tested a range of puromycin concentrations and timepoints to identify conditions that do not trigger integrated stress response to the neural progenitors. 0.1 mg/ml puromycin for up to 2 hours gave sufficient staining intensity without inducing a cell stress response. Puromycin was applied to the differentiating NPs in differentiation medium supplemented with RA and SAG at 0h timepoint. Samples were collected at the indicated timepoints for intracellular staining with anti-puromycin Alexa Fluor 488 (EDM Millipore MABE343-AF488) and SOX2-V450 conjugated antibodies.

### Global proteomics

#### Sample prep

Cells were treated with accutase to detach from cell culture plate, washed with PBS, counted and centrifuged. Cell pellets were frozen in an ethanol/dry ice bath and stored in -80oC until ready to process. Protein extracts were prepared with lysis buffer containing: 50mMTris pH8, 1% SDS, complete protease inhibitors (one tablet per 3.5 ml buffer, ROCHE 05056489001) and benzonase (250 U/ml, Pierce 88701). Protein concentration was determined using the BCA assay according to manufacturer’s instructions (Pierce 23227). To prepare peptides 100 µg of proteins per sample were acetone precipitated, washed twice with 80% acetone and solubilised in 100 mM Hepes pH 8 at 1 mg/ml protein concentration. To reduce disulfide bonds and alkylate free cysteine residues samples were treated with 10 mM DTT for 30 min followed by 20 mM Iodoacetamide for 30 min, followed again by 10 mM DTT for 30 min. Proteins were digested with LysC (1 µg LysC per 100 µg of proteins) overnight (O/N) followed by Trypsin (1.33 µg Trypsin per 100 µg of proteins) O/N. Samples were stored at -80°C until ready to proceed with mass spec.

#### Data Independent Acquisition (DIA) analysis on timsTOF Pro2

Peptides were analysed using an Evosep One LC system (EvoSep) coupled to a timsTOF Pro 2 mass spectrometer (Bruker Daltonik GmbH) using a commercial 150-mm analytical column (EV1113 ENDURANCE COLUMN, Evosep Biosystems) and an integrated Captive Spray Emitter (IonOpticks). Buffer A was 0.1% formic acid in water, Buffer B was 0.1% formic acid in acetonitrile. Data was collected using diaPASEF with 1 MS frame and 9 diaPASEF frames per cycle with an accumulation and ramp time of 100=ms, for a total cycle time of 1.07=seconds. The diaPASEF frames were separated into 3 ion mobility windows, in total covering the 400 – 1000=m/z mass range with 25=m/z-wide windows between an ion mobility range of 0.64–1.4 Vs/cm^2^. The collision energy was ramped linearly over the ion mobility range, with 20=eV applied at 0.6 Vs/cm^2^ to 59=eV at 1.6 Vs/cm^2^.

#### DIA Data Processing and Analysis

Data were analyzed using DIA-NN (version 1.8) with all settings as default except we omitted normalisation. Specifically, a maximum of one tryptic missed cleavage was allowed, with fixed modifications of N-term methionine excision and carbamidomethylation of cysteine residues. No variable modifications were selected. A *Homo sapiens and a Mus musculus* UniProt database was used for the analysis of the relevant samples. A default threshold of 1% false discovery rate was used at both the peptide and protein level. At least two unique peptides were required to identify a protein. The library-free mode of DIA-NN was used to generate precursor and fragment ions *in silico* from the UniProt database. Additionally, the programme generates a library of decoy precursors (negative controls). Retention time alignment was performed using endogenous and iRT peptides, and peak scores were calculated by comparison of peak properties between observed and reference spectra.

#### Data Analysis

The data were analysed in three samples (biological replicates) derived from three bulked samples of mouse or human NPs. Each sample was run three times on the mass-spec (three technical replicates). Protein copy numbers per cell were estimated using the proteomic ruler method^17^, and 1:1 homologs between mouse and human were retained for downstream analysis. The table with homologs was extracted from ensemble using BioMart^56^ and then manually curated to include missing gene names (Table S4). The proteins that were identified in at least 2 samples in each group (human or mouse) were used for downstream analysis. To reduce potential technical biases between mouse and human samples, we performed a global correction of protein abundances by dividing them to the size factors of the samples (total copy number of each sample per the mean of all samples). The average copy number between the non-zero technical replicates were used for differential expression, with zeros being replaced by medians intensities in each group (human and mouse). The abundances were transformed to log2 scale and differential expression was performed with R package limma^57^, using empirical Bayes method for 2 group comparison with standard t-test of the eb.fit function output^58^. The differentially expressed proteins (DEPs) were chosen based on the criteria of ≥ 2 fold change (FC) and the false discovery rate (FDR) adjusted P values for multiple-hypothesis testing with the Benjamini–Hochberg method FDR ≤ 0.05.

The pathway enrichment analysis was performed separately on mouse- or human-upregulated DEPs. We used David functional enrichment analysis^21^ with annotations from GO:BP, GO:MF, GO:CC, KEGG and Reactome databases. The background was represented by orthologous human or mouse proteins identified in mouse or human samples, respectively (Table S1). The DAVID functional annotation clustering was used to group similar and redundant terms and the top representative terms of each cluster were combined for plotting.

### Dynamic SILAC

#### Cell Culture and Stable Isotope Labeling

Mouse and human ESCs were differentiated as described above until 1^st^ and 4^th^ day of RA and SAG addition, respectively. Then, cells were collected as 0h time point, or continued the differentiation in media containing heavy arginine and lysine until collected after 3h, 8h, 24h and 48h for mouse NPs, or after 6h, 24h, 48h and 96h for human NPs in three biological replicates. For the “heavy” condition, DMEM/F12 and Neurobasal were replaced by SILAC DMEM/F12 and SILAC Neural Basal (AthenaES 0423 and 0428) supplemented with methionine and leucine. L-arginine hydrochloride (13C6, 15N4) and L-lysine hydrochloride (13C6, 15N2) (Cambridge Isotope Laboratories CNLM-539-H and CNLM-291-H) were dissolved in PBS and kept at -80°C and were used at the final concentrations of 0.549 mM and 0.648 mM, respectively, based on the concentrations in the original medium. Medium change intervals, seeding densities, coating methods and supplements of inhibitors and growth factors followed the differentiation protocols described above.

#### Sample prep

The cell pellets were lysed in 5% SDS/100mM triethylammonium bicarbonate buffer (Sigma TEAB) by probe sonication and heating. Protein concentration was measured by Pierce BCA Assay. 40 µg proteins per sample were taken, reduced by tris(2-carboxyethyl)phosphine hydrochloride solution (Sigma) and alkylated by iodoacetamide (Sigma) then purified by acetone precipitation. Proteins were digested with trypsin (0.8 µg; Sequence modified, Promega) overnight at 30°C in 50 mM ammonium bicarbonate.

#### Data Independent Acquisition (DIA) analysis on Orbitrap Eclipse Tribrid

The LC-MS/MS analysis was on the Orbitrap Eclipse Tribrid mass spectrometer coupled with a U3000 RSLCnano system (both from ThermoFisher Scientific) using DIA mode. 200 ng peptides were first loaded to the trap column (100 µm i.d. x 5 cm) and then separated on an analytical column (75 µm id x 50 cm) using a 120-min gradient from 2 - 30% acetonitrile/0.1% formic acid at a flow rate of 300 nl/min. Both trap column and analytical column were packed in-house with C18 (3 µm, Reprosil C18AQ, Dr. Maisch). MS1 scans were acquired from m/z 390 - 865 using a resolution setting at 120,000, AGC at 400,000 and maximum injection time at 50 msec. MS/MS scans were acquired in the Orbitrap in DIA mode, where the precursor mass range was set at m/z 400 – 855 with isolation windows of 16 m/z and 1 m/z overlap. The scan range was set at m/z 200 – 1800, with a resolution at 30,000, maximum injection time at 54 msec, and relative HCD collision energy at 30.

#### Data Analysis

DIA raw files were analysed using Spectronaut version 18.0 using the spectral library-free directDIA approach, searching against a human or mouse UniProtKB-proteome database (version June 2022). The Pulsar search settings were: Trypsin (specific) with maximum 2-missed cleavage sites; peptide length from 7 to 52; Carbamidomethyl (C) as fixed modification; toggle N-terminal M is true; Oxidation (M) and Acetyl (Protein N-terminus) as variable modifications. FDRs at PSM, peptide and protein level were all set to 0.01. Quantitation was based on peak areas from MS2 data, with channel 1 set for light (label-free), and channel 2 set for heavy peptides (Arg(10) and Lys(8)). All other settings were default.

Data were processed in R (v4.4.1) and RStudio (v2023.06.0+421) and mouse and human datasets were processed in parallel. The fraction of light (channel 1) compared to heavy (channel 2) chain was calculated for each replicate at each time point (L/(L+H)). Starting from 8127 and 8509 detected proteins from mouse and human NPs, respectively. Quality check filters were applied to remove proteins that had missing data, <0.85 fraction light at the 0h timepoint, <1000 total expression level over all timepoints, or unstable total expression levels across the timepoints. Instability in expression levels was defined as differences in total expression across the timepoints (Kruskal Wallis p<0.01).

Fraction of light was transformed to a log2 scale, and a linear model was fitted for each protein across the time points using all three replicates, with the y-intercept constrained to 0. Those with R2 >0.85 and sigma <0.7 were kept, and the slope of the model was used to calculate protein half-life using the following equation: 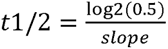

Gene set enrichment analysis on the half-lives was conducted using GSEAPreranked [v4.3.3,]^59,60^ to identify GO:BP, GP:MF, and GO:CC terms enriched in short or long half-lives. Terms were deemed significant if they had a False Discovery Rate (FDR) less than 5% (q<0.05). Revigo^61^ was used to collapse similar terms, and those with a dispensability score <0.5 were retained. Terms with fewer than 25 proteins in the dataset or that had greater than 80% overlapping genes with a term with larger normalised effect size (NES) were removed.

Homologs between mouse and human were extracted from Ensembl v112 using BioMart^56,62^, and those with 1:1 homologs were retained. Gene homologs that matched to multiple Uniprot IDs were removed. Half-life fold difference between mouse and human was calculated (human/mouse) to visualise differences in half-life between the species (Table S9).

To group the proteins by GO terms, at least one of the homologous proteins annotated with descendant terms of following were counted: CC: cytoplasm (GO:0005737), nucleus (GO:0005634), membrane (GO:0016020) and extracellular region (GO:0005576); MF: catalytic activity (GO:0003824), structural molecule activity (GO:0005198), transporter activity (GO:0005215), binding (GO:0005488), molecular adaptor activity (GO:0060090), molecular function regulator activity (GO:0098772), transcription regulator activity (GO:0140110). ATP-dependent activity (GO:0140657); For MF, only the terms with more than 100 proteins were shown, and terms with complete overlap were excluded (Table S10).

### Proteome stability measurements in neurons

AHA pulse and chase experiments were performed as in Rayon et. al 2020^4^. Cells were starved by replacing complete differentiation medium with methionine-free medium for 30 min. Next, 100mM AHA was added to the methionine-free medium for 1h. To measure protein stability, AHA pulse was removed by washing the cells once with PBS and growing the cells on complete differentiation medium for the course of the experiment. On the indicated days of differentiation and at specific time points after AHA removal, cells were processed for intracellular flow cytometry.

AHA-incorporated proteins were labeled using Click-iT™ Cell Reaction Buffer Kit (Thermo Fisher Scientific C10269) on a volume of 150 µl per sample reaction. Estimations of global proteome stability were calculated as in Rayon et al. 2020^4^. Briefly, estimations of proteome stability were obtained by normalizing each individual replicate using an initial exponential fit y(t) = B + C·exp(kt) to determine the baseline B and initial fluorescence intensity C, allowing a comparison of all the replicates together. Exponential fits of the bootstraps of the normalized ensemble were used to construct confidence intervals of the degradation rates. Error intervals reported correspond with 95% confidence intervals. At least 2 biological replicates in technical duplicates per species per time point from independent experiments were used.

### Proteasome activity measurements

Mouse and human NPs were washed with PBS once, and then scraped in proteasome assay buffer (50 mM Tris-HCl, pH7.5, 5 mM MgCl2, 0.5 mM EDTA and 10% Glycerol). Then, lysate was homogenized by passing through 27-gauge needles ten times. For the embryo extracts mouse and human embryos were dissected in cold PBS to isolate branchial to lumbar spinal cords which were subsequently homogenised in proteasome assay buffer by passing through 27-gauge needles ten times. Lysate was centrifuged at 10,000 x *g* for 10 min at 4°C and supernatant was aliquoted for storage at -80°C. 20 µg of total proteins per well of 96-well plate or 8ug of total protein per well of 384 well plate with black walls (Greiner, 655090 or 781090) were diluted in the proteasome assay buffer containing 2 mM ATP (Thermo Fisher Scientific), 1 mM dithiothreitol (Sigma) and 0.333 mg/ml Z-GGL-AMC (Cambridge Bioscience). Fluorescence was measured by PHERA star or Ensight plate reader (360 nm excitation, 430 nm emission, BMG Labtech or Perkin Elmer) every 2 min for 60 cycles at 37°C. Kinetic slopes were measured for the stable 60 min. Protein amount was estimated by BCA assay as in Vilchez et al. 2012^33^.

All animal procedures were carried out in accordance with the Animal (Scientific Procedures) Act 1986 under the Home Office project license PP8527846.

Human embryonic material (4-6 weeks of gestation) was obtained from the MRC/Wellcome-Trust (grant #006237/1) funded Human Developmental Biology Resource (HDBR57, http://www.hdbr.org) with appropriate maternal written consent and approval from the London Fulham Research Ethics Committee (23/LO/0312) and the Newcastle and North Tyneside NHS Health Authority Joint Ethics Committee (23/NE/0135). HDBR is regulated by the UK Human Tissue Authority (HTA; https://www.hta.gov.uk) and operates in accordance with the relevant HTA Codes of Practice. This work was part of project no. 200804 registered with the HDBR.

## Data and code availability

Scripts to analyse global and SILAC proteomics are both found at https://github.com/POTsnake2/Nakanoh-Stamataki_2024.

Global proteomics datasets have been deposited deposited to the PRIDE database^63^ with the project accession identifier: PXD054152.

---

**Figure S1.**
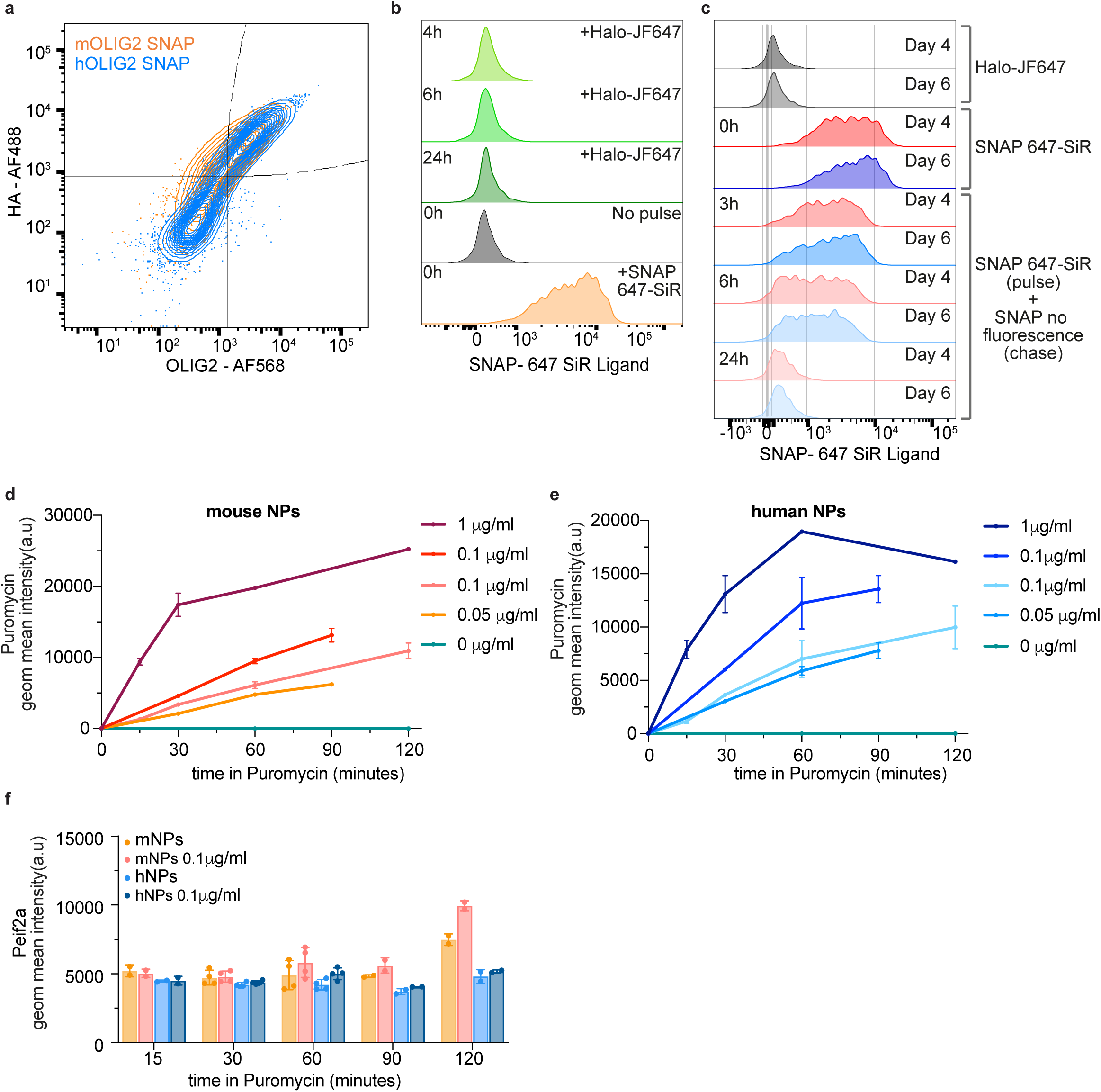
Assessment of SNAP-Tag suitability to determine protein dynamics. (a) Representative FACS plot showing co-expression of OLIG2 and HA tag in OLIG2-SNAP mouse and human NPs. (b) Representative flow cytometry analysis of 647-SiR ligand incorporation in pulse-chase experiments of human NPs at the indicated days of differentiation with HALO ligands as controls. (c) Representative flow cytometry analysis of 647-SiR ligand incorporation on day 6 (blue) in human NPs pulse-chase experiment. (d) Geometric mean ± standard deviation of fluorescent intensity of conjugated puromycin antibody in mouse NPs treated with the indicated concentrations of puromycin (n=2). (e) Geometric mean ± standard deviation of fluorescent intensity of conjugated puromycin antibody in human NPs treated with the indicated concentrations of puromycin (n=2). (f) Quantitation of phospho-EIF2A in mouse and human NPs treated or without puromycin at the indicated time points after puromycin incorporation. Data represent mean ± standard deviation (n=2 for 15, 90,120min and n=4 for 30, 60min - two experiments with different timepoints).

**Figure S2.**
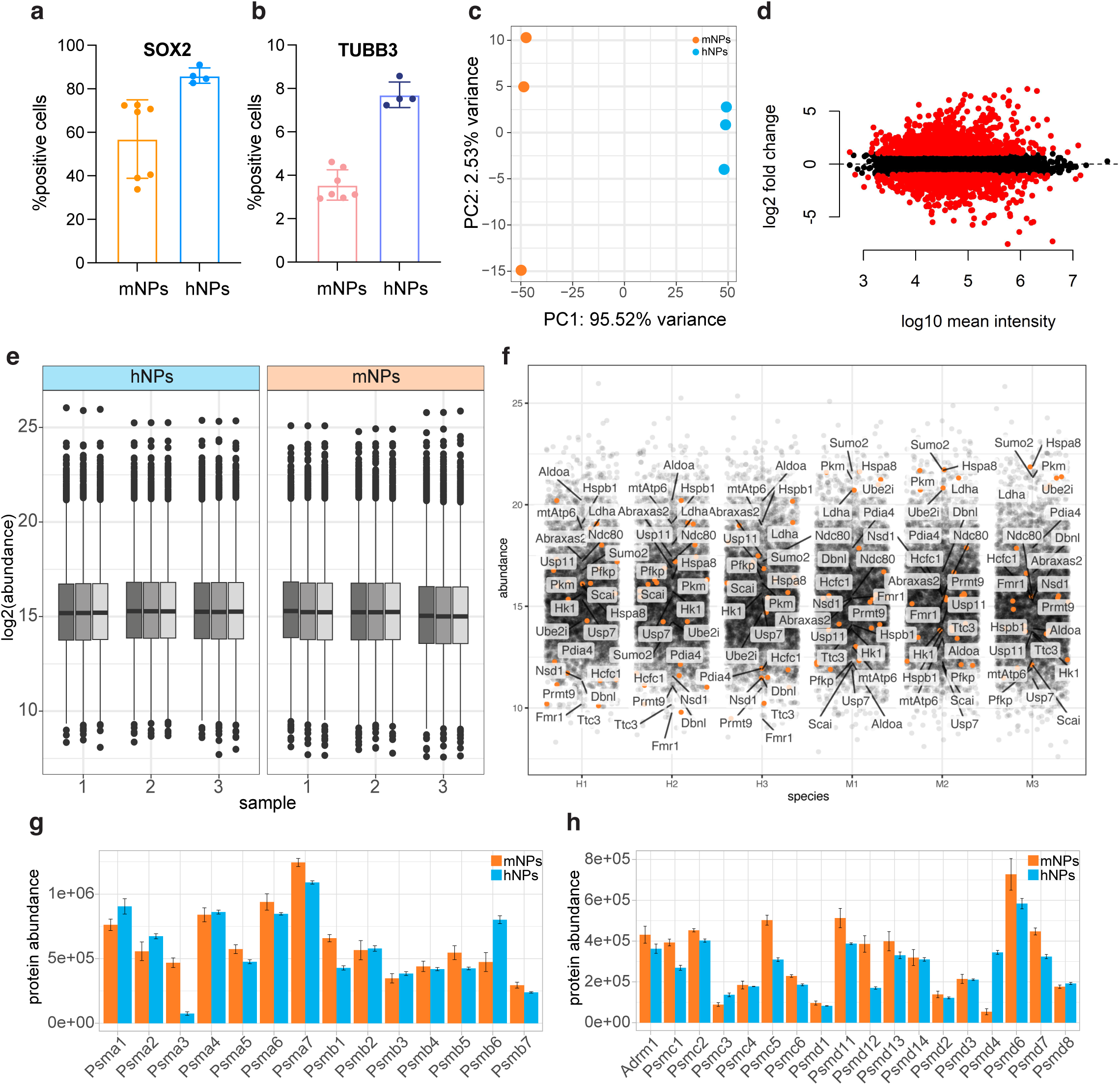
Quality control and differential expression of human and mouse proteins identified by DIA-MS in NPs. (a, b) Percentage of SOX2 (a) and TUBB3 (b) positive cells in mouse and human NP differentiations. Data represent mean ± standard deviation (mouse n=7, human n=4). (c) Principal Component Analysis plot of the mouse and human NPs used for differential protein expression. Each point represents a replicate ran in triplicate for DIA-MS. (d) MA plot displaying the logged intensity ratio (M) versus the mean logged intensities (A) of changes in protein abundances of the mouse and human NPs. The log2 of fold changes in mean human/mouse protein abundance is shown against log10 of the averaged protein abundance. Red dots indicate differentially expressed proteins. (e) Boxplots of the normalised values per technical replicate for the DIA-MS in mouse and human NPs. Each biological replicate was run in triplicate for DIA-MS. (f) Mean abundance of DEPs in mouse and human NPs across the replicates. Proteins mentioned in the main text and shown on the volcano plot are labeled. (g) Mean abundance of core particles of the proteasome in mouse and human NPs as determined by DIA-MS. Data represent mean ± standard deviation (n=3, in triplicates). (h) Mean abundance of regulatory particle subunits of the proteasome in mouse and human NPs as determined by DIA-MS. Data represent mean ± standard deviation (n=3, in triplicates).

**Figure S3.**
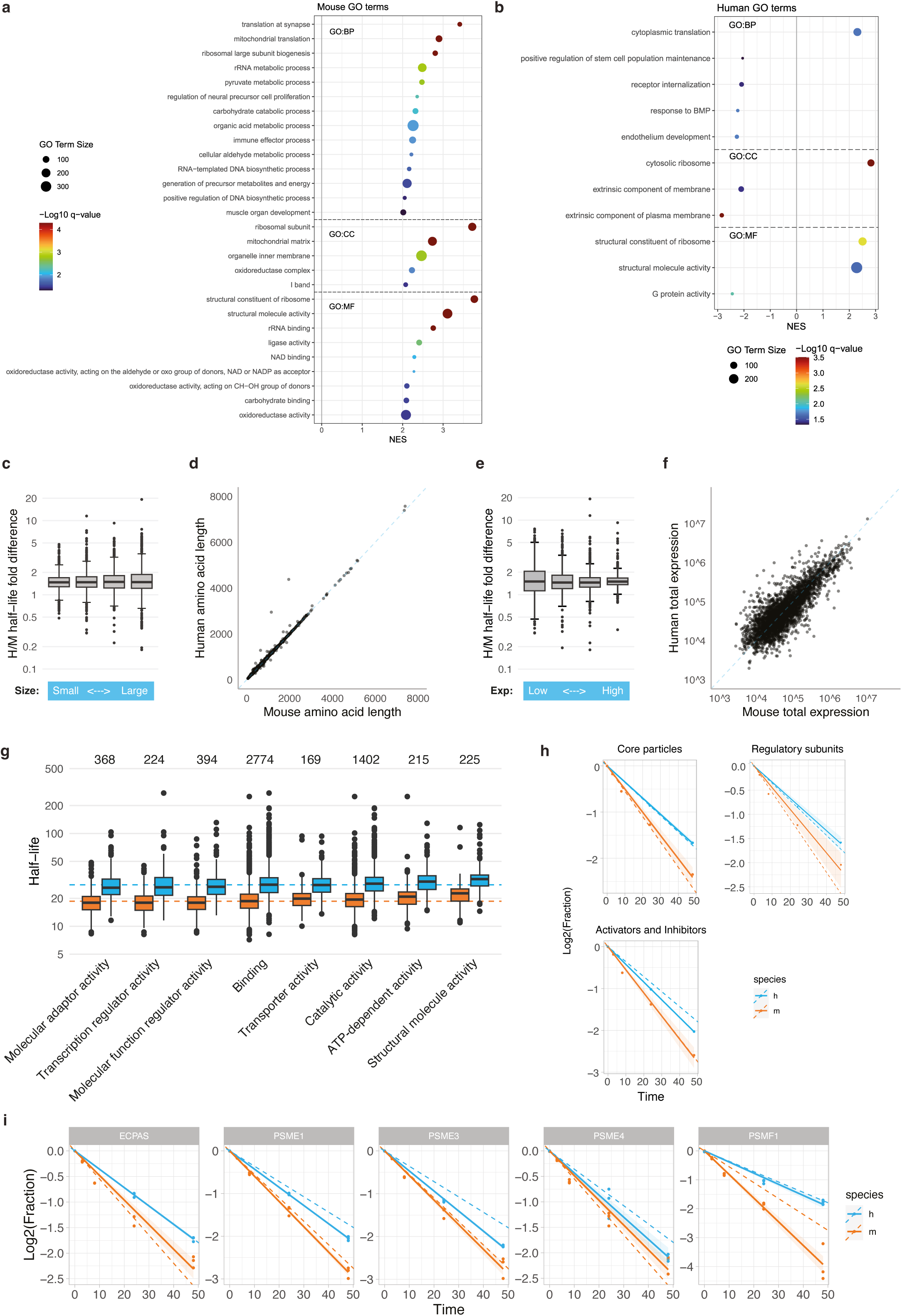
Additional data analysis of the SILAC dataset. (a, b) Bubble plot showing top enriched GO terms in mouse (a) and human (b) half-lives from GSEA Preranked analysis. Positive normalized enrichment scores (NES) indicate the enrichment in long-lived proteins, and negative NES values indicate enrichment in short-lived proteins. Size of the points depict size of the GO term, colour indicates - log10 p-value. (c) Human to mouse fold difference in half-lives in human homologous proteins partitioned in quartiles according to protein size in human, from small to large. Box-plot: Centre line is the median; box represents upper and lower quartiles; whiskers are1.5x interquartile range from the upper and lower quartiles; dots are outliers. (d) Dot plot showing amino acid sequence length of mouse to human homologues. (e) Quartiles of measured half-lives in human homologous proteins partitioned according to abundance of human proteins, from low to high. Box-plot as in (c). (f) Dot plot showing average total expression levels of mouse and human homologues. (g) Half-lives of mouse (orange) and human (blue) homologous proteins associated with 8 GO molecular function terms. Box-plot as in (c). (h) Median fraction of light over time for proteins related to proteasomal categories in the mouse and human datasets. 14 core proteasomal subunits, 16 regulatory subunits, and 5 proteasomal activators and inhibitors were taken into account. Line and shadowed areas show best linear fit with 95% confidence intervals. Dashed lines represent median protein half-lives. (i) Fraction of light over time for activators and inhibitors of the proteasome in the mouse and human NP SILAC datasets. Line and shadowed areas show best linear fit with 95% confidence intervals. Dashed lines represent median protein half-lives (n=3).

**Figure S4.**
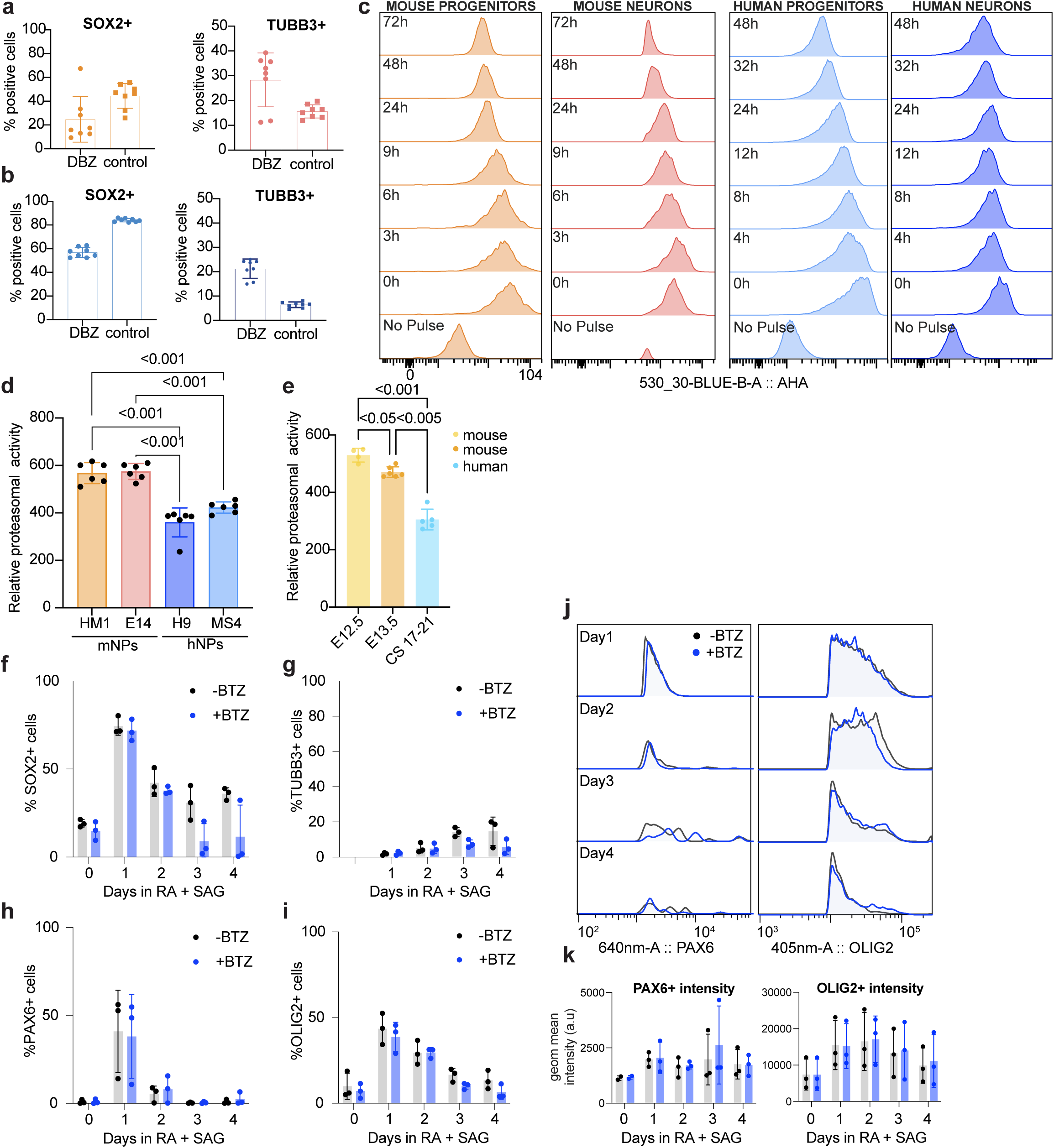
Differences in protein stability between mouse and human neural cells. (a, b) Quantifications of the proportion of neural progenitors (SOX2+) and post-mitotic neurons (TUBB3+) after treatment compared to controls in mouse (a) and human (b) differentiations. Data represent mean ± standard deviation (mouse n=2, human n=2 in duplicates). (c) Representative flow cytometry analysis of AHA incorporation in pulse-chase experiments of mouse and human NPs and neurons at the indicated time points. (d) Proteasomal activity measured as the slope of fluorescence increase from substrate digestion in mouse and human NPs, derived from two mESC lines. Data are presented as mean ± standard deviation (n = 6). Statistical analysis was performed using ordinary one-way ANOVA. (e) Proteasomal activity measured as the slope of fluorescence increase from substrate digestion in mouse and human embryonic developing spinal cords. Mouse samples are separated across stages. Data are presented as mean ± standard deviation (E12.5 (n = 4); E13.5 (n = 6); human n = 5. CS17 = 1, CS18 = 1, CS20 =2, CS21=1); Statistical analysis was performed using Brown-Forsythe ANOVA to account for unequal sample sizes. (f-i) Quantifications of the proportion of neural progenitors (SOX2+), post-mitotic neurons (TUBB3+) and early neural progenitor markers (PAX6+, OLIG2+) treated with Bortezomib (5nM BTZ) compared to controls in mouse differentiations. Data represent mean ± standard deviation (n=3). (j, k) Representative flow cytometry analysis (j) and geometric intensity (k) for PAX6+ and OLIG2+ neural progenitors treated with Bortezomib (5nM BTZ) compared to controls in mouse differentiations. Data represent mean ± standard deviation (n=3).

**Figure S5.**
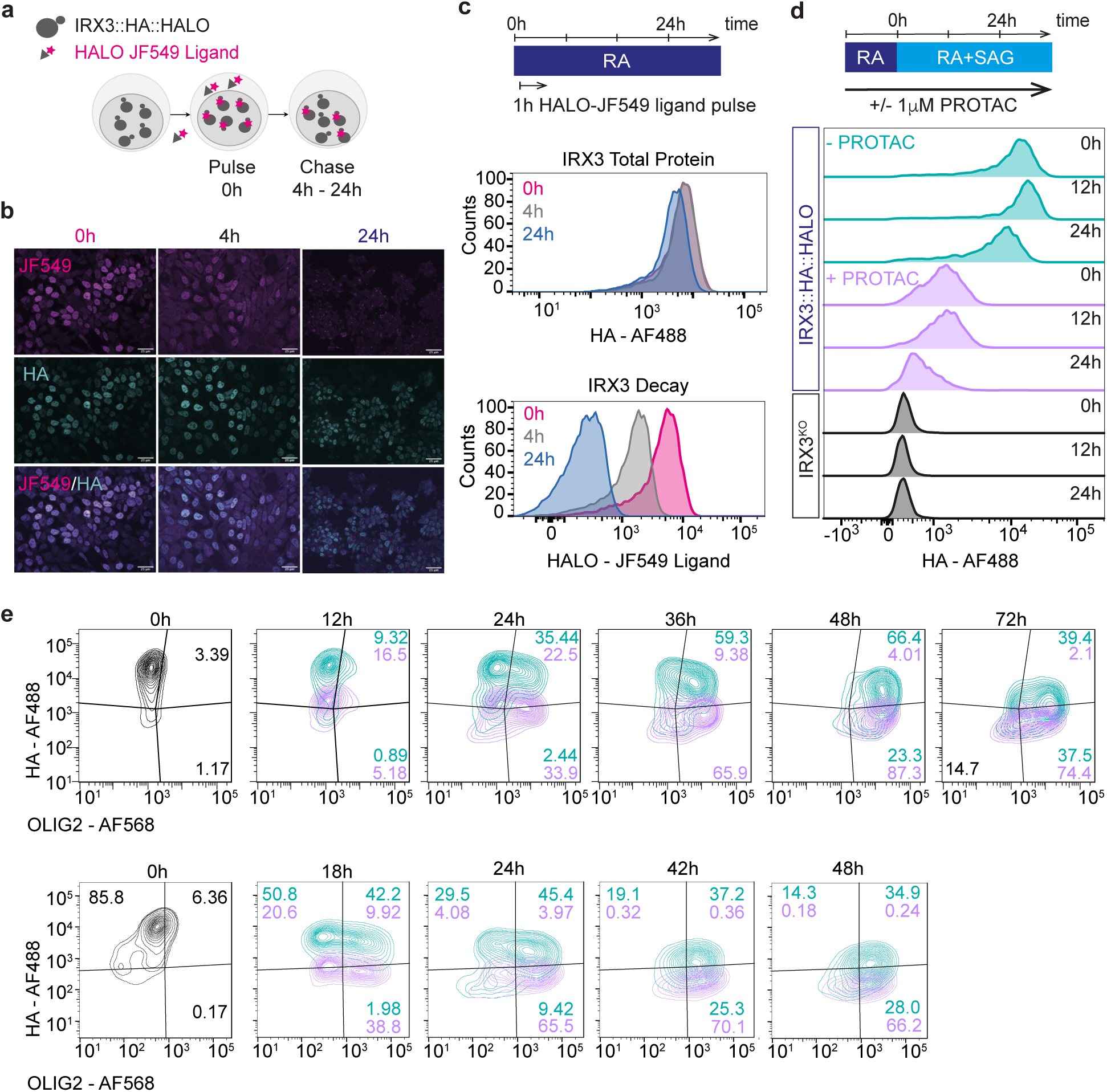
Characterization of the IRX3-HALO mESC line. (a) Diagram of pulse-chase treatment for IRX3::HALO ligand. 100nM of HALO JF549 ligand was added for 1h. Media was then replaced, and mNPs collected at the indicated time points. (b) Confocal images of IRX3 detection and decay from the pulse-chase experiments. IRX3 is detected with an antibody against the HA tag in blue throughout the time course, while the fully labelled IRX3 with the HALO JF549 ligand (magenta) at 0h shows the decay of IRX3 protein at 4h and 24h post-labelling (n=2). Scale bar 25 μm. (c) Schema indicating medium conditions in the pulse-chase labelling experiment and representative flow cytometry histograms of IRX3 expression and decay from the pulse-chase experiments. The histogram on the top corresponds to total IRX3 levels detected with an HA antibody. The lower panel corresponds to the decay of IRX3-JF549 labeled proteins at 4h and 24h post-labelling (n=2). (d) Validation of IRX3 depletion by HALO PROTAC as compared to an IRX3^KO^ cell line. Representative histograms of IRX3 expression and depletion over time upon PROTAC addition (n=2). (e) Representative contour plots of IRX3 depletion in 12h and 18h time courses. X-axis represents OLIG2 intensity and y-axis IRX3 intensity. Numbers in each quadrant indicate percentage of NPs quantified in DMSO (turquoise) or PROTAC (violet) conditions.

## ACKNOWLEDGEMENTS

We thank Miki Ebisuya for productive discussions. Rahul Samant, Ian McGough, and all members of the Briscoe lab and Rayon lab for advice and feedback. We are grateful to David Vilchez for advice and sharing the proteasome activity protocol. Thanks to Florence Wood for performing some experiments on the regimes of treatment for the IRX3-HALO line. We thank the Babraham Institute Facilities in particular the Flow Facility, Mass Spec and Bioinformatics as well as Stores, BICS, and Tech Services. The latter were instrumental when setting up the lab. We thank the Crick Science Technology Platforms for their expertise and assistance, particularly the Genomics, the Flow Cytometry, and the Proteomics STPs.

## AUTHOR CONTRIBUTIONS

T.R. and J.B. conceived the project, interpreted the data, and wrote the manuscript with input from all authors. T.R., S.N. and D.S., designed and performed experiments and data analysis. L.G.P. generated and characterized the OLIG2-SNAP and IRX3-HALO mESCs lines and designed, performed and analyzed experiments. C.A. designed and performed experiments and data analysis of IRX3-HALO mESCs. A.P. performed statistical analysis of global proteomics. L.D. performed experiments and data analysis of proteasome inhibition experiments. G.L.M.B prepared human embryo samples. M.M prepared mouse samples. S.H and M.S performed global DIA mass spec. L.Y and D.O performed SILAC mass spec analysis. H.C. and S.A. performed bioinformatic analysis of the SILAC datasets.

## FUNDING

Work in JB laboratory was supported by the Francis Crick Institute which receives its core funding from Cancer Research UK (CC001051), the UK Medical Research Council (CC001051), and Wellcome (CC001051); by the European Research Council under European Union (EU) Horizon 2020 research and innovation program grant 742138; and by UK Human Developmental Biology Initiative (Wellcome, 215116_Z_18_Z). Work in the laboratory of TR is supported by the Babraham Institute’s BBSRC Institute Strategic Programmes Grant (ISPG) [BB/Y006909/1]; the Engineering and Physical Sciences Research Council [EP/X021521/1]; the Babraham Institute’s BBSRC Core Capability Grant (CCG) [BB/CCG2210/1], and the Institute Development Grant [BB/IDG2210/1].

## COMPETING INTERESTS

The authors declare no competing or financial interests.

## Notes

### Competing Interest Statement

The authors have declared no competing interest.

### Summary of Updates

Revised version to reflect new experiments in Figure 1 and Figure 4

